# T cell receptors from CD8 regulatory T cells direct the development of natural intraepithelial lymphocytes

**DOI:** 10.64898/2026.11.11.765811

**Authors:** Samuel J.H. Murphy, Jevon Bonner, Tony Yao, Chandrani Thakur, Julia Bals, Wayne M. Yokoyama, Nicholas Borcherding, Naresha Saligrama

## Abstract

CD8 regulatory T cells (CD8 Tregs) in mice have been identified by their expression of the surface receptor family Ly49 and the transcription factor Helios. To our surprise, many Ly49+ and Helios+ T cells express neither CD4 nor CD8, and many express only CD8αα. We found that CD8 Tregs in the spleen are linked to the CD8β- natural intraepithelial lymphocytes (IELs) that reside in the small intestine. Blocking trafficking to the intestine increased splenic CD8 Treg frequency, and transferred splenic CD8 Tregs gave rise to natural IELs. Unexpectedly, Helios was dispensable for natural IEL development, arguing against its role as a shared lineage-determining transcription factor. Instead, we found that T cell receptors (TCRs) from splenic Ly49+ CD8 Tregs deterministically direct natural IEL development. Our findings suggest that CD8 Tregs may derive from the same TCR-directed developmental program that generates natural IELs and shed light on the origins of these immunoregulatory cells.

## Main Text

CD8 Tregs have been investigated for several decades^1–4^, but they remain understudied compared to the well-established CD4 Tregs that express Foxp3. Efforts to study CD8 Tregs were greatly advanced by the identification of Ly49 as a CD8 Treg marker^5^, and by the identification of Helios as a transcription factor that is important for CD8 Treg function^6^. Importantly, studies have largely focused on the ability of CD8 Tregs to control antibody responses by targeting the non-classical MHC class I molecule Qa-1b expressed by activated CD4 T cells^7^. We previously extended these observations with our discovery of a set of peptides that expanded the population of splenic Ly49+ CD8 Tregs, and that robustly suppressed autoimmunity in the setting of experimental autoimmune encephalomyelitis^8^. These peptides were presented by H2-Db and therefore showed that CD8 Tregs can also recognize peptides presented by classical MHC class I molecules. Recent observations used the genetic depletion of Ly49F-expressing cells to argue that CD8 Tregs play an important role in preventing immunopathology secondary to viral infections^9^, and it has been posited that the functional role of CD8 Tregs is to control autoreactive T cell responses^10^.

The regulatory function of CD8 Tregs has been well established in the field. However, it remains unclear how CD8 Tregs are generated, and the goal of this study was to contribute new insights on this topic. One recent study sequenced the TCRs of Qa-1b-restricted CD8 Tregs, and found a narrow TCR repertoire^11^. TCR transgenic T cells that expressed CD8 Treg TCRs displayed regulatory activity against activated CD4 T cells, reinforcing the idea that the TCR is a key determinant of CD8 Treg function. However, this study made clear that the Qa-1b-restricted TCRs make up a very small share of the total pool of Ly49+ T cell TCRs^11^.

Given our previous serendipitous discovery of classical MHC class I-restricted CD8 Tregs, we wanted to use as unbiased a definition of CD8 Tregs as possible, to avoid inadvertently missing important aspects of CD8 Treg biology. Using only Ly49 as a marker of our cells of interest, here we sought to better understand the origins and developmental trajectories of Ly49+ T cells. We were surprised to find that many splenic Ly49+ T cells do not express CD8. In addition to cells that lacked TCR co-receptors entirely, there was a prominent population of Ly49+ T cells that expressed only CD8αα, and they were exclusively Helios+. The existence of CD8αα+Ly49+ T cells in the spleen suggested a connection to the natural intraepithelial lymphocytes (natural IELs, also referred to as unconventional IELs) that are found in the small intestine, many of which express CD8αα (some lack expression of CD8 altogether, so they are defined as CD4-CD8β-IELs)^12–15^. We investigated the relationship between CD8 Tregs and natural IELs, and the importance of Helios for natural IEL development. We focused on the TCR as a key link between these populations, and we found that TCRs from splenic CD8 Tregs give rise to natural IELs. Taken together, our observations argue that splenic CD8 Tregs and natural IELs are part of a shared developmental lineage.

## Results

### Many Ly49+ and Helios+ T cells do not express a TCR co-receptor

Consistent with earlier reports^16^, we noticed that Ly49+CD8+ T cells bear less CD8 on their surface than other CD8+ T cells (Fig. 1a,b). We were surprised to find that many Ly49+ T cells lack CD8 expression altogether (Fig. 1c). When we examined TCRβ+CD4-T cells, to exclude CD4 T cells but include all other αβTCR+ cells, we found that nearly 40% of all Ly49+ T cells were CD8-negative (Fig. 1d; Extended Data Fig. 1a,b). This called into question whether CD8 is a key feature of the so-called “CD8” Tregs that have been defined on the basis of Ly49 expression^5^. To begin to evaluate whether CD8 expression is associated with a distinct sub-population of Ly49+ T cells, we examined as unbiased a population of splenic Ly49+ T cells as possible by sorting CD3+ cells that express Ly49 from the spleens of C57BL6/J mice and performing single cell RNA sequencing (scRNAseq).

**Figure 1:**
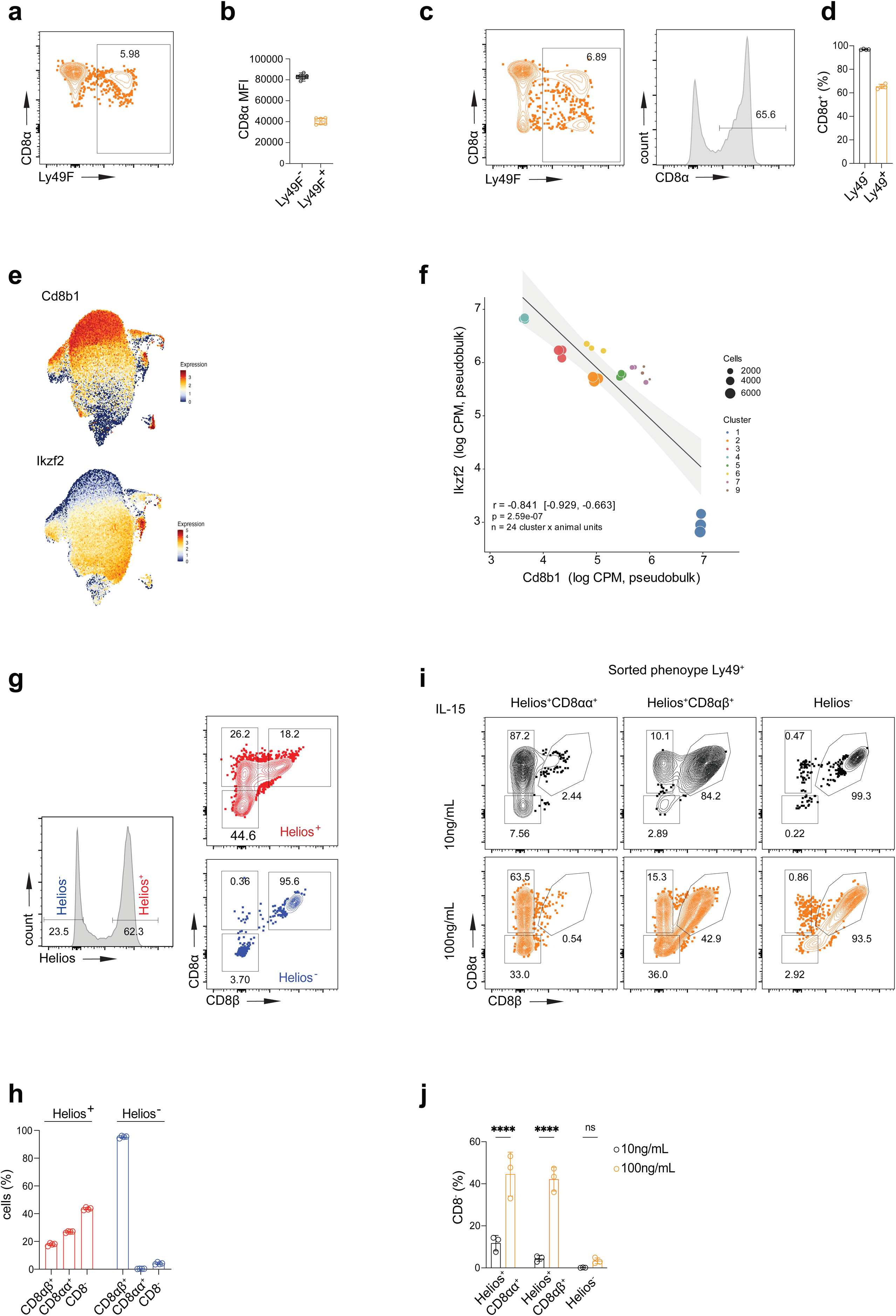
Many Ly49+ and Helios+ T cells do not express a TCR co-receptor. **a,** Representative flow plot showing Ly49F and CD8α expression among splenic cells pre-gated as CD8α+ T cells. **b,** Quantification of CD8α mean fluorescence intensity (MFI) for Ly49F+ and Ly49F-CD8+ T cells from **a** (n=6 mice). **c,** Representative flow plot showing Ly49F and CD8α expression among splenic cells pre-gated as TCRβ+CD4-T cells. Subsequent gating for CD8α+ cells among total splenic Ly49+ T cells is shown (see Extended Data Fig. 1 for gating of total Ly49+ T cells). **d,** Quantification of CD8α frequency among Ly49+ and Ly49-splenic TCRβ+CD4-T cells (n=4 mice). (**e,f),** scRNAseq was performed on CD3+Ly49+ splenic T cells from C57BL/6J mice (n=3 mice). **e,** Feature plots showing gene expression of *Cd8b1* and *Ikzf2* overlaid on a UMAP of all sequenced Ly49+ T cells. **f,** Pearson correlation of *Ikzf2* and *Cd8b1*, with each cluster treated as a pseudobulk population. **g,** Representative flow plots showing Helios expression among Ly49+ T cells from Helios-GFP reporter mice, and expression of CD8α and CD8β among Helios+ and Helios-Ly49+ T cells. Frequency of each gated subset is quantified in **h** (n=4 mice). **(i,j)**, Splenic CD8α+Ly49+ T cells were divided based on Helios expression, and Helios+Ly49+ T cells were further divided based on CD8β expression. Sorted cells were split and cultured in either low-dose (10ng/mL) or high-dose (100ng/mL) IL-15. **i,** Representative flow plots showing expression of CD8α and CD8β after 11 days in culture. **j,** Quantification of frequency of CD8-cells in each of the conditions for the sorted populations (n=3 mice). Statistical significance was calculated using a one-way ANOVA followed by Šídák’s multiple comparisons test; ns P≥0.05, **** P<0.0001. Data are presented as mean values ± SD.

As expected, the sorted cells were heterogeneous, and included iNKT cells (*Plzf*+) and a small number of terminal effector (*Klrg1*+) CD8 T cells (Extended Data Fig. 2a,b). We considered the other populations to be putative CD8 Tregs, and these included cells undergoing interferon signaling and proliferation (Extended Data Fig. 2a,b). The most striking finding was the relationship between CD8 and Helios expression in the sequenced Ly49+ T cells. Putative “CD8” Tregs seemed to be divided into either conventional-appearing CD8αβ+ T cells that lacked Helios expression (encoded by *Ikzf2*), or cells that were high for Helios but lacked CD8β expression, with some intermediate transcriptional states also present (Fig. 1e). By treating the clusters as bulk populations, the strong anti-correlation between *Ikzf2* and *Cd8b1* expression was readily apparent (Fig. 1f).

Using Helios-GFP reporter mice^17^ and flow cytometry, we confirmed that Ly49+ T cells could be clearly divided into Helios+ and Helios-subsets. The Ly49+Helios-cells were almost uniformly CD8αβ+ T cells, while few Ly49+Helios+ cells were CD8αβ+ (Fig. 1g,h). Instead, the plurality of Ly49+Helios+ cells did not express CD8 on their surface at all. Of note, there was also a sizable population of Helios+Ly49+ T cells that expressed CD8α without CD8β, presumably in the form of CD8αα homodimers. To test the durability of CD8 expression on the surface of these different populations, we divided CD8+Ly49+ T cells based on their expression of Helios and CD8β (Fig. 1i). Borrowing from the NK cell field, where high-dose IL-15 (100ng/mL) can activate NK cells and low-dose IL-15 (10ng/mL) can maintain NK cells in vitro^18^, we assessed whether IL-15 might have an effect on CD8 expression. When maintained with low-dose IL-15, the sorted Ly49+ T cell populations retained their respective sorted phenotypes (Fig. 1i, top row). When cultured instead with high-dose IL-15, many of the Helios+ cells (whether sorted as CD8αβ+ or CD8αα+) lost CD8 expression (Fig. 1i, bottom row). This did not occur to an appreciable degree for the Ly49+Helios-cells (Fig. 1j). Thus, CD8 seemed not to be a stable surface marker for Ly49+Helios+ T cells. Based on these findings, we focused subsequent analysis on TCRβ+CD4-T cells, to avoid excluding any relevant cells that lack CD8 expression.

### Splenic Helios+CD122+ T cells share several features with natural IELs of the small intestine

Among Ly49+ T cells, about 30% of the Helios+ subset express the α4β7 integrin, while hardly any of the Helios-subset express α4β7 (Fig. 2a,b; Extended Data Fig. 3a). Along with their expression of the CD8αα homodimer (Fig. 1g), this suggested a connection to the CD8β-natural IELs that reside in the small intestine^12–15^. Indeed, all CD8β-natural IELs express Helios (Fig. 2c,d). Type A thymic IEL precursors (IELp)^19^ also uniformly express Helios (Extended Data Fig. 3b,c). We hypothesized that Helios+CD122+ T cells in the spleen are connected to this intraepithelial T cell lineage, and we tested this hypothesis in several ways.

**Figure 2:**
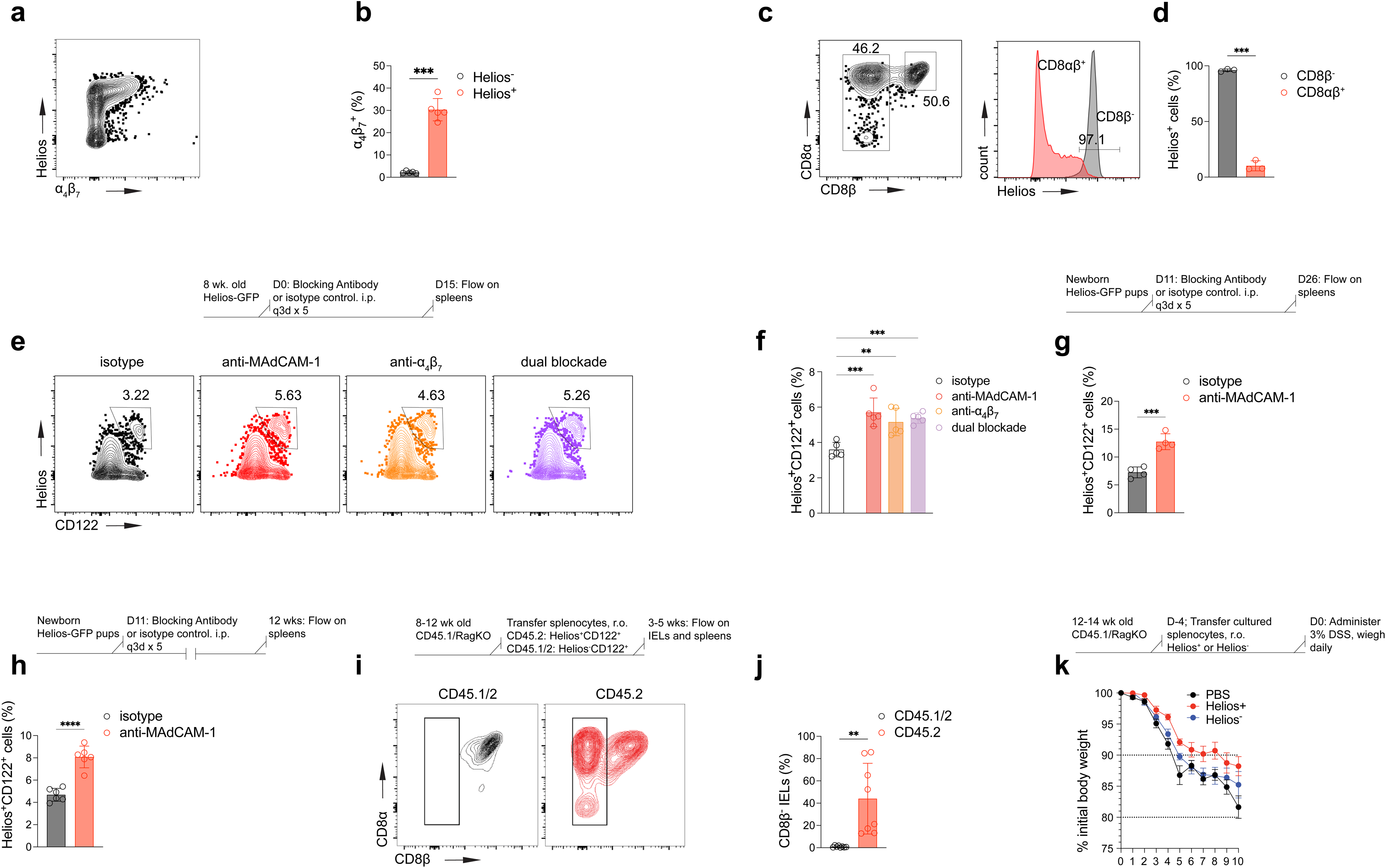
Splenic Helios+CD122+ T cells share several features with natural IELs of the small intestine. **a,** Representative flow plot showing integrin α4β7 and Helios expression among total splenic Ly49+ T cells (TCRβ+CD4-). **b,** Quantification of frequency of α4β7 expression among Helios+ and Helios-Ly49+ T cells (n=5 mice). **c,** Representative flow plots showing CD8α and CD8β expression on pre-gated TCRβ+CD4-IELs from the small intestine (left), and Helios expression among CD8αβ+ and CD8β-IELs (right). **d,** Quantification of Helios expression among CD8αβ+ and CD8β-IELs (n=3 mice). **e,** Antibodies against α4β7 and/or MAdCAM-1 were used to disrupt trafficking to the intestine in adult mice. Top: schematic illustrating antibody treatment. Bottom: representative flow plots showing CD122 and Helios expression among splenic TCRβ+CD4-T cells for the different treatment groups. **f,** Quantification of frequency of Helios+CD122+ cells among splenic TCRβ+CD4-T cells (n=5 mice per group). **g,** Antibody against MAdCAM-1 was used to disrupt trafficking to the intestine in neonatal mice. Top: schematic illustrating antibody treatment. Bottom: quantification of frequency of Helios+CD122+ cells among splenic TCRβ+CD4-T cells (n=4 mice per group). **h,** Antibody against MAdCAM-1 was used to disrupt trafficking to the intestine in neonatal mice, and mice were left untreated for an additional 8-9 weeks. Top: schematic illustrating antibody treatment. Bottom: quantification of frequency of Helios+CD122+ cells among splenic TCRβ+CD4-T cells (n=6 mice per group). **i,** Congenically distinct Helios+CD122+ (CD45.2) and Helios-CD122+ (CD45.1/2) T cells were transferred into RagKO (CD45.1) recipients. Top: experimental schematic. Bottom: representative flow plot showing CD8α and CD8β expression among congenically distinct cells recovered from the small intestine IEL compartment. **j,** Quantification of frequency of CD8β-IELs from the different donor populations. Data are pooled from two independent experiments (n=8 mice in total). **k,** Helios+CD122+ and Helios-CD122+ T cells were separately sorted and expanded in IL-15 for 7 days before being transferred into RagKO recipients. Recipient mice were subjected to DSS for 10 days. Top: experimental schematic. Bottom: weight loss curves. Data are pooled from two independent experiments (n=10 mice per group in total). Statistical significance was calculated using a mixed-effects model. The overall treatment effect was significant (P=0.0151). Multiple comparison testing was performed for each treatment compared to PBS, and Dunnett’s multiple comparisons test was used. Only the comparison of PBS vs. Helios+ gave statistically significant differences. For the other panels, statistical significance was calculated using Welch’s T test (**b**), a one-way ANOVA followed by Dunnett’s multiple comparisons test (**e**), unpaired T tests (**f,g**), and a paired T test (**i**); ** P<0.01, *** P<0.001, **** P<0.0001. Data are presented as mean values ± SD.

First, we disrupted the ability of immune cells to traffic to the gut, using antibodies that block either α4β7 or its ligand MAdCAM-1^20,21^. Blocking gut trafficking with either antibody in adult animals increased the frequency of splenic Helios+CD122+ T cells, and there was no apparent additive effect of blocking both molecules (Fig. 2e,f). Previously, a thymocyte time-stamping approach revealed that the natural IEL compartment is preferentially seeded during a narrow window early in life^22^. We found that blocking gut homing during this window resulted in a dramatic increase in splenic Helios+CD122+ T cells (Fig. 2g). Surprisingly, this effect persisted long after the end of antibody treatment; mice treated with MAdCAM-1 blockade as neonates continued to have increased frequencies of Helios+CD122+ T cells in their spleens 8 weeks later (Fig. 2h), despite an increased frequency of α4β7 expression among the splenic Helios+CD122+ T cells after neonatal MAdCAM-1 blockade (Extended Data Fig. 3d). This suggests that T cells which are prevented from following their natural trafficking pattern to the intestine appear as CD8 Treg-phenotype cells (Helios+CD122+) in the spleen and may persist in that state for an extended period.

Next, we tested the ability of splenic T cells to give rise to natural IELs. We hypothesized that only CD8 Tregs, defined by Helios expression, would adopt the natural IEL phenotype after homing to the intestine. We isolated Helios+CD122+ T cells (CD45.2) and comparator Helios-CD122+ T cells (CD45.1/2) from the spleens of donor mice and co-transferred the cells into congenically distinct RagKO recipients (Fig. 2i). We chose Helios-CD122+ T cells as the comparator population because IL-15 is required for natural IEL generation^23,24^, and CD122 is part of the receptor for IL-15^25^. Though the frequency in the IEL compartment varied, many of the Helios+CD122+ T cells that homed to the small intestinal epithelium became natural IELs (CD8β-). The co-transferred Helios-CD122+ T cells, in contrast, gave rise exclusively to CD8αβ+ IELs (Fig. 2i,j). Thus, among IL-15-responsive splenocytes, only Helios+ T cells have the potential to become natural IELs.

It has been reported that natural IELs and their thymic precursors can protect RagKO;IL2rgKO mice from chemically induced colitis^24^. To determine if splenic Helios+CD122+ T cells could similarly protect mice from colitis, we isolated splenic Helios+CD122+ T cells and comparator Helios-CD122+ T cells and expanded them with IL-15 in vitro for 7 days before transferring them into RagKO recipients. Mice that received Helios+CD122+ splenocytes had slightly reduced weight loss compared to PBS controls, but Helios-CD122+ T cells did not confer protection (Fig. 2k). Collectively, these observations show that many features of natural IELs are shared by Helios+CD122+ splenic T cells.

### Helios is dispensable for natural IELs

The high level of Helios expression by both splenic CD8 Tregs and natural IELs (Fig. 1g,2c) made Helios a strong candidate for a lineage-determining transcription factor that might unite these two populations. To test the importance of Helios, we started by deleting Helios in CD8 T cells using Helios floxed^26^ (referred to as Helios^f/f^ here) and E8i-Cre^27^ (referred to as Cd8^Cre^ here) mice. Likely owing to their lack of CD8 expression, many Ly49+ T cells in the spleen did not demonstrate Cd8^Cre^ activity (Extended Data Fig. 4a), and Helios^f/f^Cd8^Cre^ mice showed only a modest reduction in splenic Helios+CD122+ T cells (Extended Data Fig. 4b). Therefore, we focused our analysis on natural IELs of the small intestine. Cd8^Cre^ very efficiently deleted Helios among CD8αα+ IELs (Fig. 3a).

**Figure 3:**
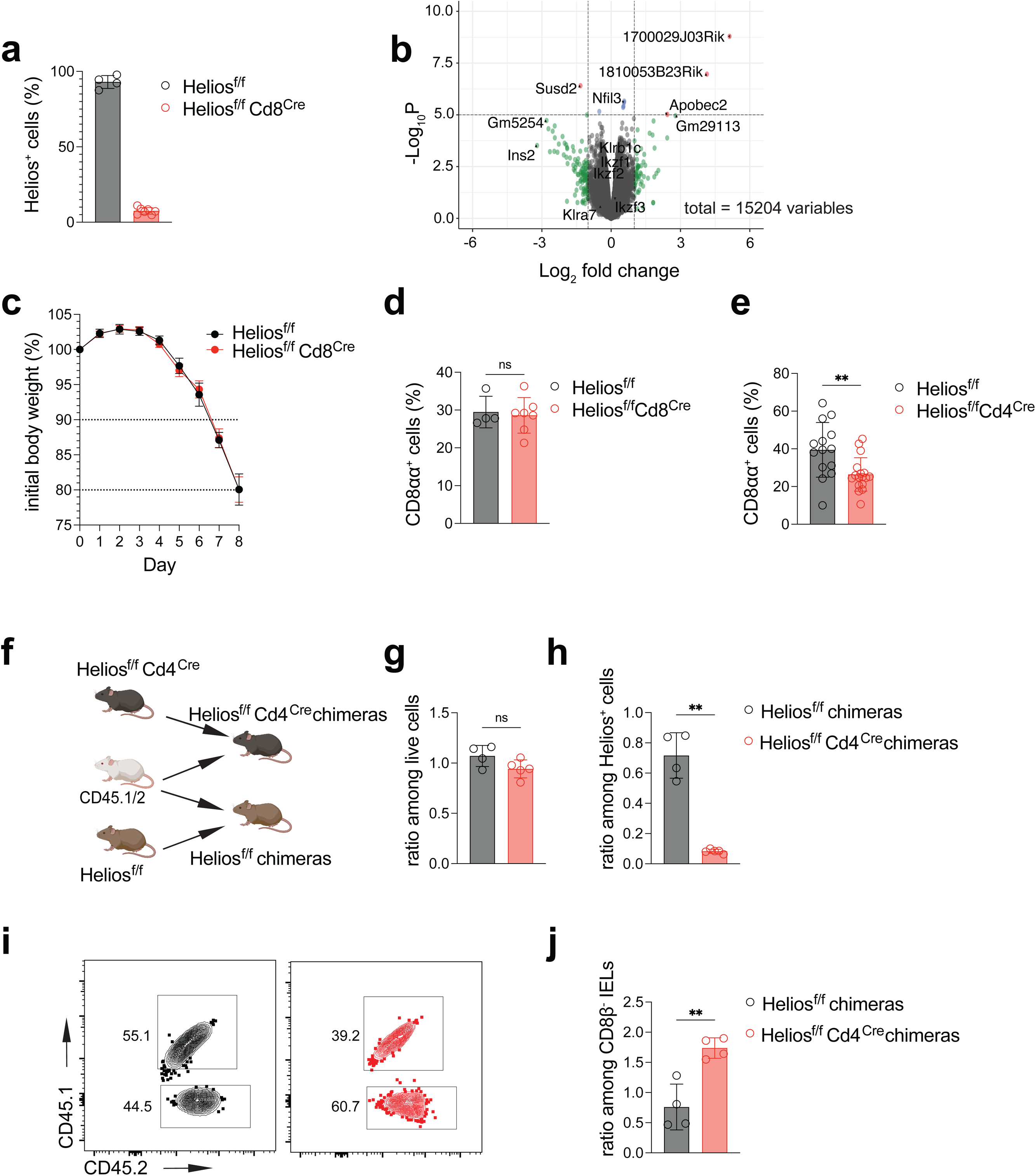
Helios is dispensable for natural IELs. **a,** Frequency of Helios+ cells among CD8αα+ IELs from Helios^f/f^ and Helios^f/f^Cd8^Cre^ mice (n=4, Helios^f/f^; n=7, Helios^f/f^Cd8^Cre^). **b,** Bulk RNAseq was performed on sorted CD8αα+ IELs from Helios^f/f^ and Helios^f/f^Cd8^Cre^ mice (n=3 per genotype). A volcano plot shows the results of differential gene expression analysis comparing Cre+ and Cre-samples. **c,** Weight loss curves for mice subjected to 8 days of DSS treatment (n=4, Helios^f/f^; n=7, Helios^f/f^Cd8^Cre^). **d,** Frequency of CD8αα+ IELs among TCRβ+CD4-IELs in mice from **c**. **e,** Frequency of CD8αα+ IELs among TCRβ+CD4-IELs from Helios^f/f^ and Helios^f/f^Cd4^Cre^ mice. Data are pooled from 4 independent experiments (n=14, Helios^f/f^; n=17, Helios^f/f^Cd4^Cre^). **(f-h),** Mixed bone marrow chimeras were used to evaluate the cell-intrinsic dependency of natural IELs on Helios (n=4, Helios^f/f^ chimeras; n=5, Helios^f/f^Cd4^Cre^ chimeras). **f,** Schematic of mixed bone marrow chimera generation. CD45.1/2 comparator marrow was mixed with either Helios^f/f^ or Helios^f/f^Cd4^Cre^ marrow to generate the mixed chimeras. **g,** Ratio of CD45.2:CD45.1/2 cells showing chimerism among total splenic cells. **h,** Ratio of CD45.2:CD45.1/2 cells showing chimerism among splenic Helios+ T cells. **i,** Representative flow plots showing chimerism among CD8β-IELs. **j,** Ratio of CD45.2:CD45.1/2 cells showing chimerism among CD8β-IELs. Statistical significance was calculated using Welch’s T test (**d,e,g,h,j**); ns P≥0.05, ** P<0.01. Data are presented as mean values ± SD.

Contrary to our expectations, we could find no discernible effect of Helios deletion on natural IELs. We performed bulk RNAseq on sorted CD8αα+ IELs, and we detected essentially no significantly differentially expressed genes (DEGs) when we compared natural IELs from Helios^f/f^Cd8^Cre^ and littermate Helios^f/f^ control mice (Fig. 3b). *Ikzf2* did not appear as a DEG, but we confirmed that Helios was deleted as expected, because reads that aligned to the floxed exon of *Ikzf2* (exon 8) were essentially absent from the Cre+ samples (Extended Data Fig. 4c). Next, we used DSS colitis as a model to test the functional effect of loss of Helios among IELs. Weight loss curves were nearly identical among Cre+ and Cre-animals (Fig. 3c), and there was also no difference in the frequency of natural IELs after DSS treatment (Fig. 3d). We concluded that Helios is dispensable for the maintenance and function of natural IELs.

Previous experiments that established a role for Helios among CD8 Tregs had perturbed Helios early in T cell development, by either germline deletion or via Cd4-Cre-mediated recombination at the double positive (DP) stage in the thymus^6^. The E8i-Cre used here (Fig. 3a-d; Extended Data Fig. 4a-c) is known to be active starting later, at the CD8 single positive stage in the thymus^28,29^. Therefore, we considered that Helios might only be important early during the thymic development of natural IELs, as shown recently for BCL6^30^. We crossed Helios^f/f^ and Cd4^Cre^ mice, and we found that Cd4^Cre^ led to efficient deletion of Helios among natural IELs (Extended Data Fig. 4d). Helios^f/f^Cd4^Cre^ mice showed a slight reduction in the frequency of natural IELs and thymic IELp (Fig. 3e, Extended Data Fig. 4e). Splenic Ly49+ T cells also demonstrated some changes, with a reduced frequency of CD8αα+ cells in the absence of Helios (Extended Data Fig. 4f). These results were consistent with our hypothesis that Helios may play a role early in the development of CD8 Tregs and natural IELs, but the effect sizes were small. We were also concerned that our results could be confounded by the effect of loss of Helios in CD4 Tregs^6^.

We considered that in a competitive setting, Helios deficient IELs could be at a disadvantage, and this might uncover a more pronounced effect of loss of Helios. We generated mixed bone marrow chimeras, where either Helios^f/f^Cd4^Cre^ or control Helios^f/f^ marrow was co-transplanted with CD45.1/2 competitor marrow into irradiated CD45.1 recipients (Fig. 3f). In the spleen, we confirmed that overall engraftment was comparable regardless of genotype (Fig. 3g). Helios was efficiently deleted among cells derived from the Helios^f/f^Cd4^Cre^ marrow, reflected by very low chimerism among the Helios+ cells in the Helios^f/f^Cd4^Cre^ chimeras (Fig. 3h). We noted that the chimerism among Helios+ splenocytes in the control Helios^f/f^ chimeras was also slightly less than 1:1; this could possibly be due to some disadvantage attributable to the floxed allele, but the difference was small. Overall, this confirmed that the mixed chimeras could be used to evaluate the necessity of Helios for natural IEL development.

In the IEL compartment, the results were contrary to our expectations. The Helios-deficient cells outcompeted their competitor cells and made up a greater share of the natural IEL compartment in the Helios^f/f^Cd4^Cre^ chimeras, while the Helios sufficient cells were roughly equal in proportion to the competitor cells in the control Helios^f/f^ chimeras (Fig. 3i,j). There were no clear differences in the frequency of thymic IELp in the mixed chimeras (Extended Data Fig. 4g). This suggests that the reduced frequency of natural IELs that we observed in Helios^f/f^Cd4^Cre^ mice (Fig. 3e) may be due to cell-extrinsic effects, and if anything, that these effects overcome an advantage conferred by cell-intrinsic loss of Helios (Fig. 3i,j). Our overarching conclusion is that Helios is not required for the generation of natural IELs and appears to be dispensable for their function. While Helios may be a useful marker of cells of this lineage, it does not appear to be a key unifying factor that connects CD8 Tregs and natural IELs.

### The splenic Ly49+ T cell TCR repertoire includes IEL-associated sequence motifs

We sought to identify a different feature that would support or reject the idea of a unified lineage shared by CD8 Tregs and natural IELs. Previous work established that the development of natural IELs is directed by their TCRs^31,32^, so we examined paired TCR sequencing from our scRNAseq dataset to investigate whether the TCR might lend further insight into the connection between CD8 Tregs and natural IELs. The overall splenic Ly49+ T cell TCR repertoire did not differ substantially from bulk splenic T cells. Rank-abundance plots revealed the absence of any dominant, highly expanded clones, and comparable evenness compared to the TCR repertoire of bulk splenic CD8 T cells taken from ImmGenT (Fig. 4a). TCR gene usage was diverse and similar to that of total splenic CD8 T cells (Fig. 4b), though we did observe a somewhat preferential usage of TRBV12-2 and less usage of TRBV13-2 among Ly49+ T cells (Fig. 4b).

**Figure 4:**
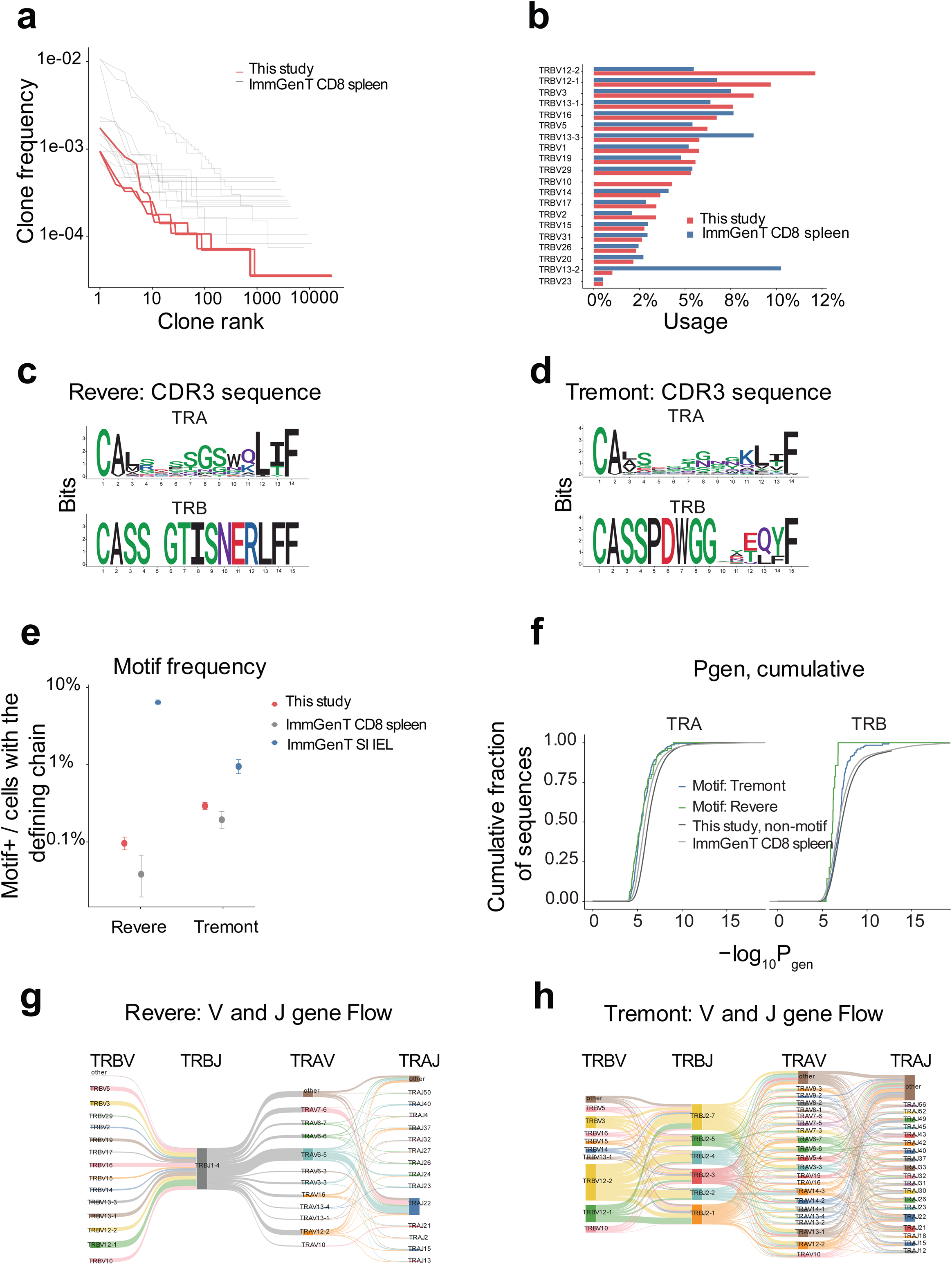
The splenic Ly49+ T cell TCR repertoire includes IEL-associated sequence motifs. **a,** Rank-abundance plot showing clone frequency for TCR clones from our splenic Ly49+ T cell TCR data (“This study”, n=3 mice), with data taken from ImmGenT splenic CD8 T cells shown as a comparator. **b,** TRBV gene usage among our splenic Ly49+ T cell TCR data (“This study”), with data taken from ImmGenT splenic CD8 T cells shown as a comparator. **c,d,** Logos plots showing definitions for two previously identified IEL-associated CDR3β sequence motifs. We defined the Revere motif (**c**) by CDR3β amino acids “CASSxGTISNERLFF” and the use of TRBJ1-4, with no constraint placed on CDR3α. We defined the Tremont motif (**d**) by CDR3β amino acids “CASSPDWGGx”, with no constraint placed on either TRBJ or on CDR3α. **e,** Frequency of motif-bearing cells in our splenic Ly49+ T cell TCR data (“This study”), ImmGenT data for splenic CD8 T cells, and ImmGenT data for IELs from the small intestine. **f,** Probability of generation (Pgen) was calculated for TRB chains that bear either a Revere or a Tremont motif (right), and their paired TRA chains (left). Included for comparison are Pgen results for non-motif splenic Ly49+ T cell TCRs from our data, and for splenic CD8 T cell TCRs from ImmGenT. **g,h,** Gene-flow plots that show the V- and J-gene usage for paired TCRs that contain either a Revere (**g**) or Tremont (**h**) motif. Each ribbon represents the proportion of TCR clones across the motif.

A previous report sequenced small intestinal IELs and identified CDR3β sequence motifs that were highly enriched among natural IELs^33^. Informed by this work, we specifically looked for “Revere”-motif TCRs that contain a core CDR3β ‘xGTI’ motif and that use TRBJ1-4, and for “Tremont”-motif TCRs that contain a core CDR3β ‘PDWGGx’ motif (Fig. 4c,d). We identified many instances of TCRs bearing these IEL-associated CDR3β motifs in our dataset (Fig. 4e). We found that Revere- and Tremont-motif TCRs have a slightly higher probability of generation (Pgen) than total splenic T cells, which suggests that they are more easily generated by VDJ recombination and may appear at a higher frequency owing to this ease of generation (Fig. 4f). Even so, Revere and Tremont sequence motifs appeared at higher frequencies in our Ly49+ T cell TCR data compared to total splenic CD8 T cells (Fig. 4e). The frequency of these motifs was highest among TCRs from the small intestine IEL compartment (data from ImmGenT). Revere- and Tremont-motif TCRs in our dataset were clearly not instances of clonal expansion, because TCRs with identical CDR3β motifs could: (1) be found in separate mice, (2) demonstrated diverse V gene usage, and (3) paired with different TCRα chains (Extended Data Fig. 5). Gene-flow plots illustrate the many different TRBV genes that combine with TRBJ1-4 to generate Revere-motif TCRs (Fig. 4g). Tremont-motif TCRs are, by definition, even more diverse, with many different TRBV and J gene pairings (Fig. 4h). TCRs bearing either of the CDR3β motifs paired with diverse TCRα chains (Fig. 4g,h).

A more recent study investigated TCR sequences from IELs in the mammary glands, and identified a CDR3α amino acid sequence that they called the “Newbury” motif^34^. We found TCRs containing Newbury motifs in our splenic Ly49+ T cell TCR data (Extended Data Fig. 6a,c). We also found what appeared to be a variant of the Newbury motif (Extended Data Fig. 6b,c). We were wary that these CDR3α motifs may primarily reflect the recombination of certain TRAV and TRAJ genes, so we chose to focus our efforts on TCRs bearing the Revere and Tremont CDR3β motifs.

The presence of TCRs bearing IEL-associated motifs among our Ly49+ T cell dataset was strongly suggestive of a link to natural IELs. Nevertheless, these TCRs were derived from splenic T cells, without any selection criteria that related to their ability to seed the intestine. Furthermore, motif-bearing TCRs are highly diverse, and it was not clear that the presence of a certain CDR3β motif alone could identify TCRs that direct natural IEL development. With these caveats in mind, we wanted to experimentally test the ability of splenic Ly49+ T cell TCRs bearing Revere and Tremont motifs to give rise to natural IELs.

### Splenic Ly49+ T cell TCRs drive the development of natural IELs

TCR retrogenic mice—bone marrow chimeras, where retrovirally-encoded TCR genes are introduced into donor bone marrow—enable one to test the ability of a TCR to determine the fate of developing T cells. It has been appreciated for some time, however, that premature expression of TCRα by traditional retrogenic (and TCR transgenic) approaches results in the aberrant generation of “lineage-confused” or “gamma/delta-like” T cells^31,35^. Many of these cells go on to reside in the intestinal epithelium as natural IELs, making this approach unsuitable for testing the ability of a TCR to direct natural IEL development. To prevent developing thymocytes from following this abnormal developmental trajectory, pioneering work introduced a floxed-stop cassette (lox-STOP-lox) upstream of the retrovirally-encoded TCRα^35^. Cd4^Cre^ was used to “time” the expression of TCRα to the DP stage, to more closely mimic normal thymic development.

To date, timed TCR retrogenic mice have been generated using donor mice with a fixed TCRβ chain, so that only TCRα must be supplied in the timed retroviral construct. One recent report described the use of a timed TCRαβ expression construct for in vitro thymocyte cultures^36^.

We developed a variation on an established method so we could generate timed TCR retrogenic mice without a fixed TCRβ chain and without recombination of the endogenous TCR locus, using Cd4^Cre^RagKO mice as bone marrow donors. We inserted an irrelevant TCRβ chain (taken from the OT-I TCR) into the floxed-stop cassette, and included a 2A-linked paired TCRαβ in place of TCRα (Fig. 5a). Thus, transduced RagKO thymocytes use the floxed TCRβ chain to pass beta selection at the double negative stage of thymic development. At the DP stage, Cd4^Cre^ removes the STOP cassette containing the irrelevant TCRβ chain, enabling the expression of the paired TCRαβ of interest. This system successfully enabled the expression of different TCRs, confirmed by staining for retrovirally encoded Vβ chains (Extended Data Fig. 7a).

**Figure 5:**
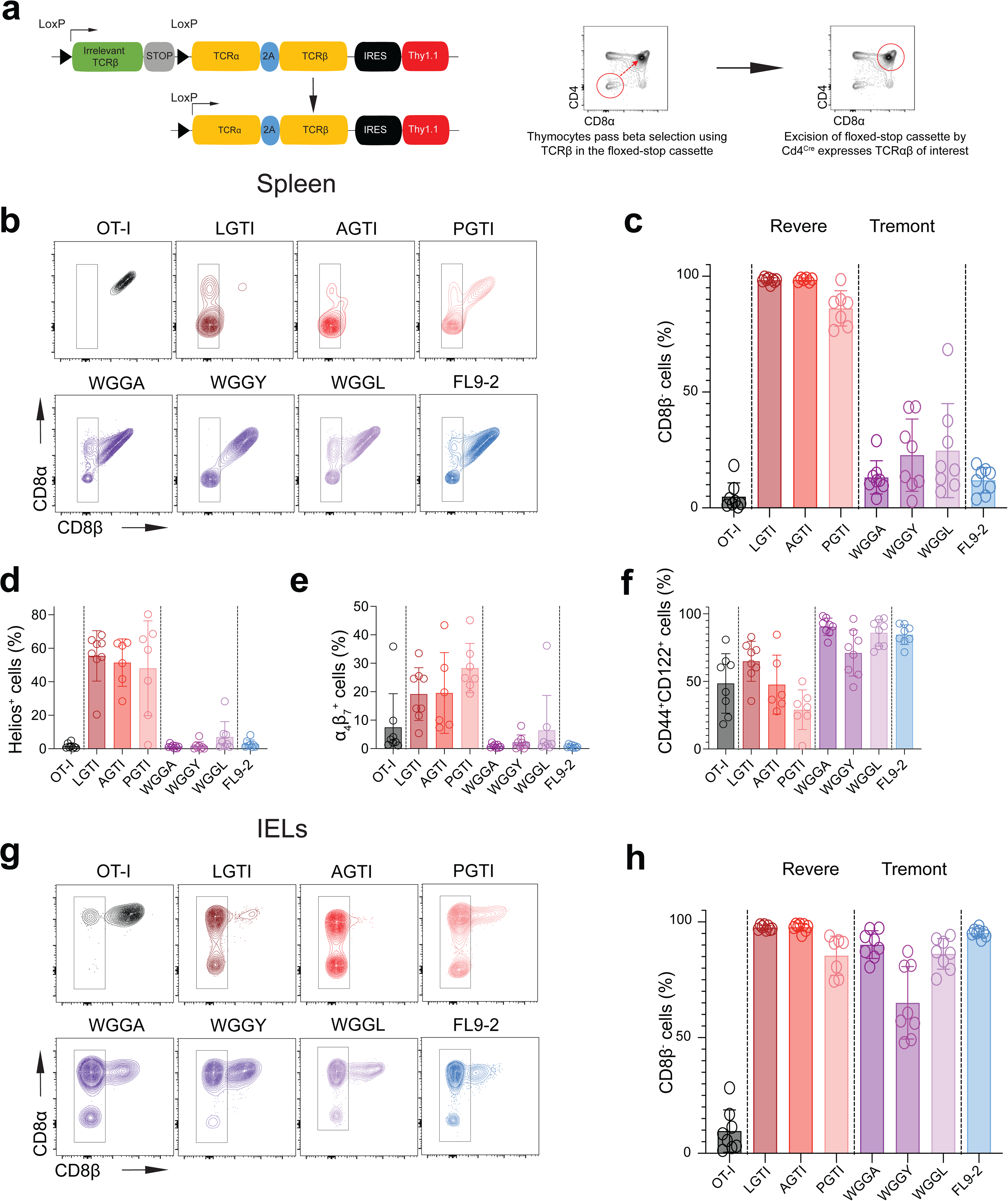
Splenic Ly49+ T cell TCRs drive the development of natural IELs. **a,** Schematic depicting the timed TCRrg construct. **(b-h),** TCRrg mice were generated for 8 different TCRs. All TCRrg mice were generated as “low frequency” retrogenics, using wildtype filler marrow to provide competitor T cells and therefore a more homeostatic developmental environment. Data are pooled from two independent cohorts of TCRrg mice per TCR (n=7 or 8 mice per TCR). **b,** Representative flow plots showing CD8β and CD8α expression among splenic TCRrg cells. **c,** Quantification of CD8β-frequency among splenic TCRrg cells. **d,** Quantification of Helios+ frequency among splenic TCRrg cells. **e,** Quantification of α4β7+ frequency among splenic TCRrg cells. **f,** Quantification of CD44+CD122+ frequency among splenic TCRrg cells. **g,** Representative flow plots showing CD8β and CD8α expression among TCRrg cells in the IEL compartment. **h,** Quantification of CD8β- frequency among TCRrg cells in the IEL compartment.

To confirm the ability of our system to give rise to expected T cell developmental trajectories, we used the OT-I TCR—known to give rise to conventional CD8 T cells—as a control. We also selected 3 TCRs bearing the Revere motif and 3 TCRs bearing the Tremont motif to test their ability to give rise to natural IELs. For the Revere and Tremont TCRs, the 3 chosen TCRs each used a different TRBV gene, and the Tremont motif TCRs also each used different TRBJ genes (Extended Data Fig. 7c).

To start, we examined the fate of TCRrg T cells in the spleen (Extended Data Fig. 7b). As expected, TCRrg T cells expressing the OT-I TCR were exclusively CD8αβ+ splenocytes that lacked Helios expression (Fig. 5b-d). Interestingly, the two IEL-associated motifs appeared to direct different splenic phenotypes. Cells bearing Revere-motif TCRs almost uniformly lacked co-receptor expression in the spleen (Fig. 5b,c). Cells bearing Tremont-motif TCRs, on the other hand, were predominantly CD8αβ+, with only a few cells either lacking a co-receptor or expressing only CD8α (Fig. 5b,c). There was also considerable Helios expression among cells bearing Revere-motif TCRs, but minimal Helios was detectable among cells bearing Tremont-motif TCRs (Fig. 5d). We detected α4β7 integrin expression on some of the Revere-motif expressing cells (Fig. 5e). CD44 and CD122 were expressed by a large number of cells regardless of the TCR they expressed, perhaps owing to the lymphopenic environment after irradiation (Fig. 5f).

Based on the results from the spleen, we expected that only the Revere-motif TCRs were likely to direct natural IEL development. We were surprised to find that in fact, both Revere and Tremont motif TCRs gave rise predominantly to natural IELs in the small intestine. As expected, the OT-I TCR gave rise exclusively to CD8αβ+ IELs (Fig. 5g). All of the IEL-associated TCRs gave rise predominantly to CD8β-natural IELs. This confirmed our hypothesis that motif-bearing TCRs derived from splenic Ly49+ T cells can direct the development of natural IELs. Unfortunately, we could not detect enough TCRrg cells in the thymus to analyze developing TCRrg thymocytes (Extended Data Fig. 7d).

We wanted to use our timed TCRrg system to make one additional connection between CD8 Tregs and natural IELs, by testing a “bona fide” CD8 Treg TCR. A recent study sequenced Qa-1b-restricted CD8 Treg TCRs, and generated TCR transgenic mice. These TCR transgenic T cells were able to suppress activated CD4 T cells^11^. Therefore, we chose to express this established CD8 Treg TCR (“FL9-2”) in our retrogenic system. In the spleen, the FL9-2 TCR demonstrated a phenotype like the Tremont-motif TCRs: predominantly CD8αβ+, with no detectable Helios or α4β7 expression (Fig. 5b-e). In the IEL compartment, however, the FL9-2 cells were almost exclusively CD8β-IELs (Fig. 5g,h). Thus, a confirmed CD8 Treg TCR also directs the development of natural IELs.

## Discussion

Our overarching argument is that splenic CD8 Tregs and natural IELs are likely to share the same developmental origins. This idea is supported most strongly by data from our TCRrg system. We found that TCRs from splenic Ly49+ T cells—and one verified CD8 Treg TCR from the literature—deterministically direct the development of natural IELs (Fig. 5g,h). Coupled with results from gut-homing blockade (Fig. 2e-h) and splenocyte transfers (Fig. 2i,j) that were undertaken in a polyclonal setting, these data argue that CD8 Tregs found in the spleen are part of a T cell lineage that is shared with the natural IELs of the small intestine. In the future, it needs to be clarified the extent to which splenic CD8 Tregs are in transit to the intestine, are recirculating from the intestine, or—as is suggested by the persistence of increased CD8 Treg frequency long after neonatal gut-homing blockade (Fig. 2h)—are cells which remain in circulation after previously failing to enter the intestine.

The finding that natural IELs are preferentially seeded early in life^22^ informed our neonatal integrin blockade experiments, but it also raises questions about the role of the neonatal environment in shaping CD8 Treg development. Another study used time-stamping to investigate splenic CD8 T cells that are generated early in life and found that cell-intrinsic factors distinguish T cells with early-life origins from adult-derived T cells^37^. Future work should explore whether CD8 Tregs from early life are shaped by the neonatal environment and whether they differ from CD8 Tregs generated in adulthood.

Previous efforts to understand thymic IELp were critical for the conceptualization of our study. Foundational studies revealed the ability of individual TCRs to selectively drive natural IEL fate choice in the thymus^31,32^. These studies challenged us to develop a TCRrg system to rigorously test our hypothesis that CD8 Treg TCRs direct natural IEL development. One of the limitations of our study is that we were unable to analyze TCRrg cells in the thymi of our TCRrg mice (Extended Data Fig. 7d), and as a result, we do not know the thymic phenotype directed by the TCRs we studied. For example, we are interested to know whether a given TCR preferentially directs the development of either type A or type B IELp^19^, and what the implications might be for the function of those TCRrg cells.

The identification of recurrent TCR motifs among natural IELs by Schattgen et al.^33^ allowed us to prioritize splenic Ly49+ T cell TCRs from our dataset to test. Given the well-established role of thymic agonist selection in directing the natural IEL fate^31,32,38–40^, we suppose that the TCRs we tested encounter agonist ligands in the thymus, and this directs their development into natural IELs. The role of antigen recognition in other tissues—whether the spleen or the intestine—remains unclear, especially considering that natural IELs appear to persist in the absence of surface TCR expression^22^. Regardless, the existence of recurrent CDR3β motifs among TCRs that direct natural IEL development raises the possibility that these diverse TCRs may recognize similar antigens. The identification of endogenous ligands for the TCRs we studied would enable the dissection of natural IEL and CD8 Treg development in greater detail than has been possible to date.

We arrived at our investigation of natural IELs by taking an unbiased approach to investigate CD8 Tregs, using Ly49 and Helios as molecular identifiers to guide our efforts. Unexpectedly, our approach raised several incongruities with previous results in the CD8 Treg literature. First, despite being defined as CD8 Tregs, we found that many Ly49+ T cells lack CD8 expression. It is possible that CD8-Ly49+ T cells may simply be unrelated to CD8 Tregs. This seems unlikely, given that cells which lack CD8 are highly enriched for Helios, the other defining molecule of CD8 Tregs (Fig. 1e-j). We found that Tremont-motif-bearing TCRs and the FL9-2 TCR could give rise to predominantly CD8αβ+ splenic T cells, while also yielding an appreciable population of CD8β-splenic T cells (Fig. 5b,c). Thus, the same TCR can give rise to splenic T cells with and without CD8 expression, consistent with a shared origin for these populations.

Nevertheless, the contribution of CD8 expression to CD8 Treg function should be considered in light of studies of mice bearing the D227K mutation in Qa-1b, which specifically disrupts binding by CD8 and thereby prevents the activation of CD8 T cells by peptide presented on Qa-1b^41^. Cantor and colleagues elegantly established that the D227K mutation abrogates the suppressive effects of CD8 Tregs^7^, which suggests that CD8 is required for CD8 Treg function. What, then, are the implications for “CD8” Tregs that lack CD8 expression? One possibility is that a subset of CD8 Tregs do not depend on antigen recognition to carry out their functions and instead behave in an innate-like manner (perhaps by responding to IL-15). Alternatively, certain CD8 Treg TCRs may be activated by their cognate antigen in the absence of the CD8 co-receptor. It has been shown that agonist selection directs thymic IELp development^31,32,38–40^, therefore it is plausible that a subset of high-affinity TCRs that are agonistically selected may be activated by their ligands in the periphery in the absence of co-receptor activity. It will be important to investigate the exact settings in which the co-receptor function of CD8 is necessary for CD8 Tregs to carry out their functions. One point is clear: by gating on CD8+ cells for their analyses, previous studies excluded many Helios+ “CD8” Tregs from investigation. Therefore, it will be critical to examine the Ly49+ and Helios+ T cells that lack a co-receptor to provide clarity on this topic.

Second, it was previously argued that Helios is a key molecule for CD8 Treg function, but our results argue that Helios is not required for either the development or the function of natural IELs. In the absence of Helios, we detected minimal gene expression changes among CD8αα+ IELs, and we did not detect functional consequences of the loss of Helios (Fig. 3b,c). In mixed bone marrow chimeras, Helios-deficient IELs readily competed against Helios-sufficient IELs (Fig. 3i,j). How can these results be squared with the previously reported importance of Helios for CD8 Treg function? The most likely answer lies in the complexity of the Ikaros family of transcription factors, which includes five members that can variably form homo- and heterodimers^42,43^. Recent work found that Aiolos is important for controlling the activation of natural IELs^44^. It seems likely that loss of Helios in natural IELs is compensated by Aiolos. Why Aiolos seems to be unable to compensate for Helios among splenic CD8 Tregs is not clear, but the interplay between different Ikaros family members in cells found in different tissues is a topic for future investigation.

One additional possibility to consider is that Helios may subtly shape the TCR repertoire of developing CD8 Tregs and natural IELs. Helios could influence the threshold for agonist selection in the thymus: in the absence of Helios, certain TCRs may fail to be appropriately selected, and other TCRs may instead fall within the threshold for agonist selection. This could explain why natural IELs appear to develop without issue in the absence of Helios, but systems where TCR specificity is important (via recognition of peptides presented by Qa-1b) reveal Helios’ significance. Since we did not evaluate the TCR repertoire upon loss of Helios, we cannot rule out such a model.

What defines a CD8 regulatory T cell? This question illustrates a tension that lies at the center of this study. We have presented evidence to support the notion that CD8 Tregs may be part of a broader lineage that includes natural IELs, but efforts to demarcate the boundaries of this lineage are complicated by recent descriptions of novel T cell populations that may or may not be related. One example is the recently described GPR15-guided regulatory CD8 “T_IGR_” cells^45^. These cells were shown to depend on GPR15 to traffic to the colon and suppress inflammation, and they also express Helios and Ly49. Another example is the progeny of “clonally evicted” thymocytes, some of which express Ly49 and Helios and appear to possess regulatory properties^46^. IELs that populate the mammary gland during lactation are yet another related population that expresses Helios^47^, especially owing to the presence of Revere motifs among their TCRs^34^. Most confusingly, intra-tumoural T cells named “innate-like T cell killers” (ILTCKs) are also candidates for this lineage: ILTCKs express Ly49 and Helios^48^, respond readily to IL-15, and their TCRs direct the development of natural IELs^49^. Though they operate in a very different context, ILTCKs can be said to be “regulatory” in the sense that they maintain homeostasis by killing malignant cells.

These disparate studies appear to describe overlapping T cell populations, but how these populations correspond to one another is unclear. A pressing need in the field is to determine the extent to which these results describe the same lineage of T cells in different contexts, or instead T cells with distinct developmental and functional features. With the present study, we have made an initial effort to draw out a previously unappreciated connection between CD8 Tregs and natural IELs. Our work and others^8,11,31,32,46,49^ have shown that approaches grounded in the TCR represent the best path towards developing a comprehensive understanding of this T cell lineage, which is necessary if basic discoveries about these cells are to be converted into translational applications.

## Extended Data Figure Legends

**Extended Data Figure 1:**
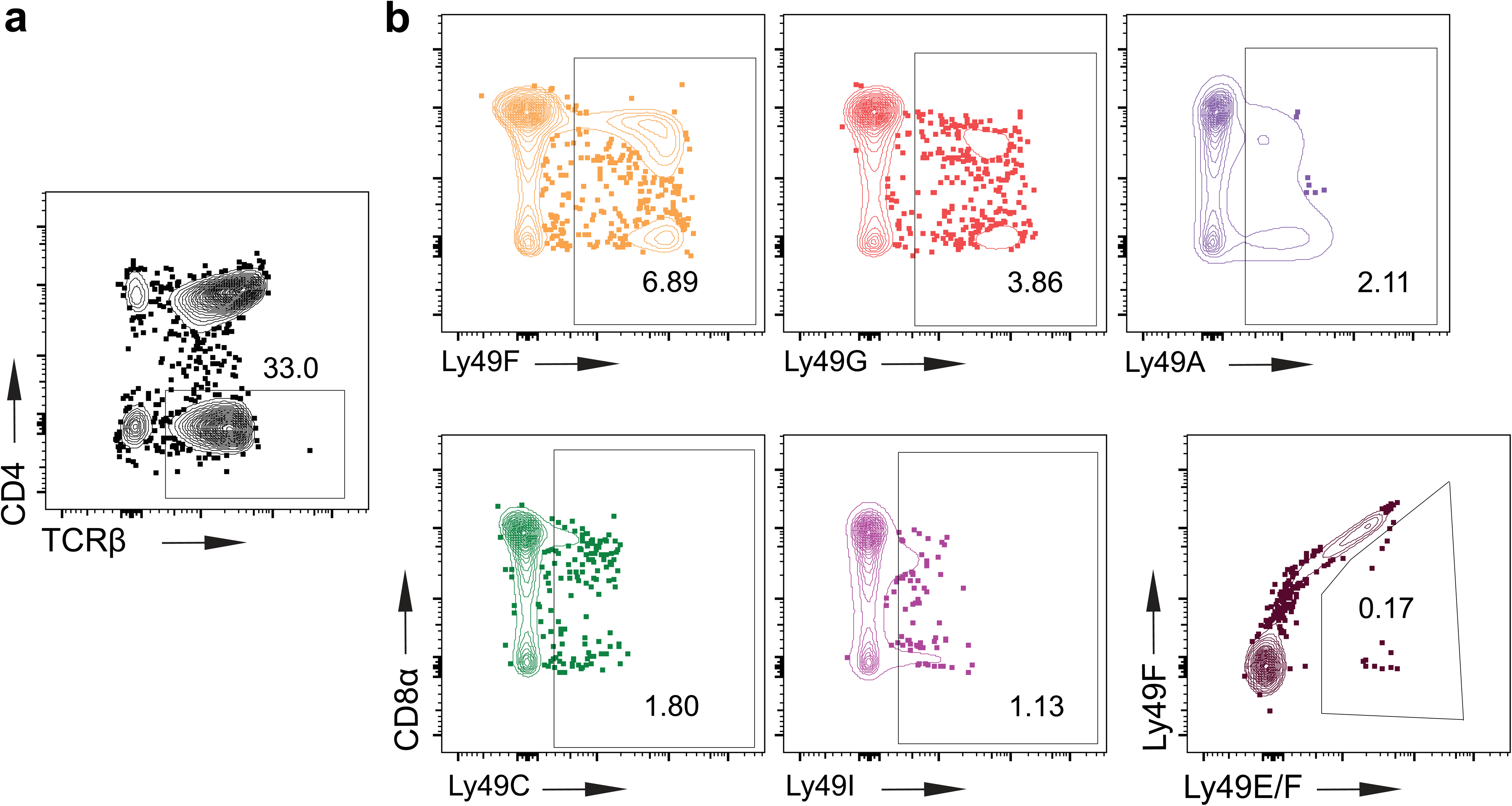
Gating strategy to identify all splenic Ly49+ T cells. **a,** Splenocytes were pre-gated as lymphocytes, singlets, live cells, CD3+ T cells, and were then gated as TCRβ+CD4- T cells. **b,** Among TCRβ+CD4- T cells, Ly49+ T cells were identified using monoclonal antibodies that are specific for each inhibitory Ly49 family member (Ly49E, for which the only monoclonal antibody available cross-reacts with Ly49F, being the only exception). Each Ly49 family member was gated separately, as shown, and the gated cells were combined for subsequent analysis as total splenic Ly49+ T cells.

**Extended Data Figure 2:**
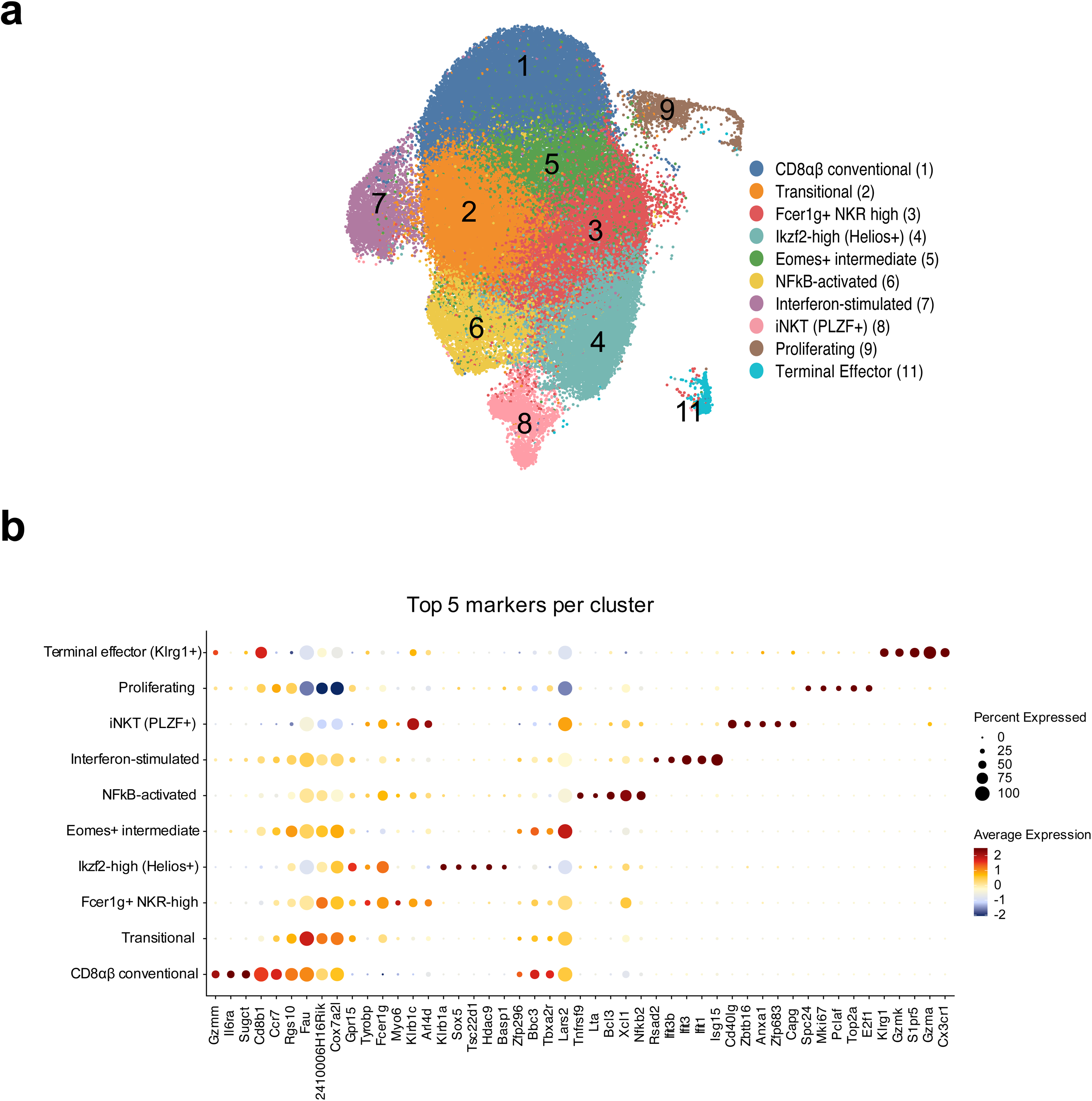
UMAP and cluster-defining genes for scRNAseq data from splenic Ly49+ T cells. **a,** UMAP of scRNAseq data generated from CD3+Ly49+ splenic T cells. **b,** Expression levels of the top 5 marker genes for each cluster are shown.

**Extended Data Figure 3:**
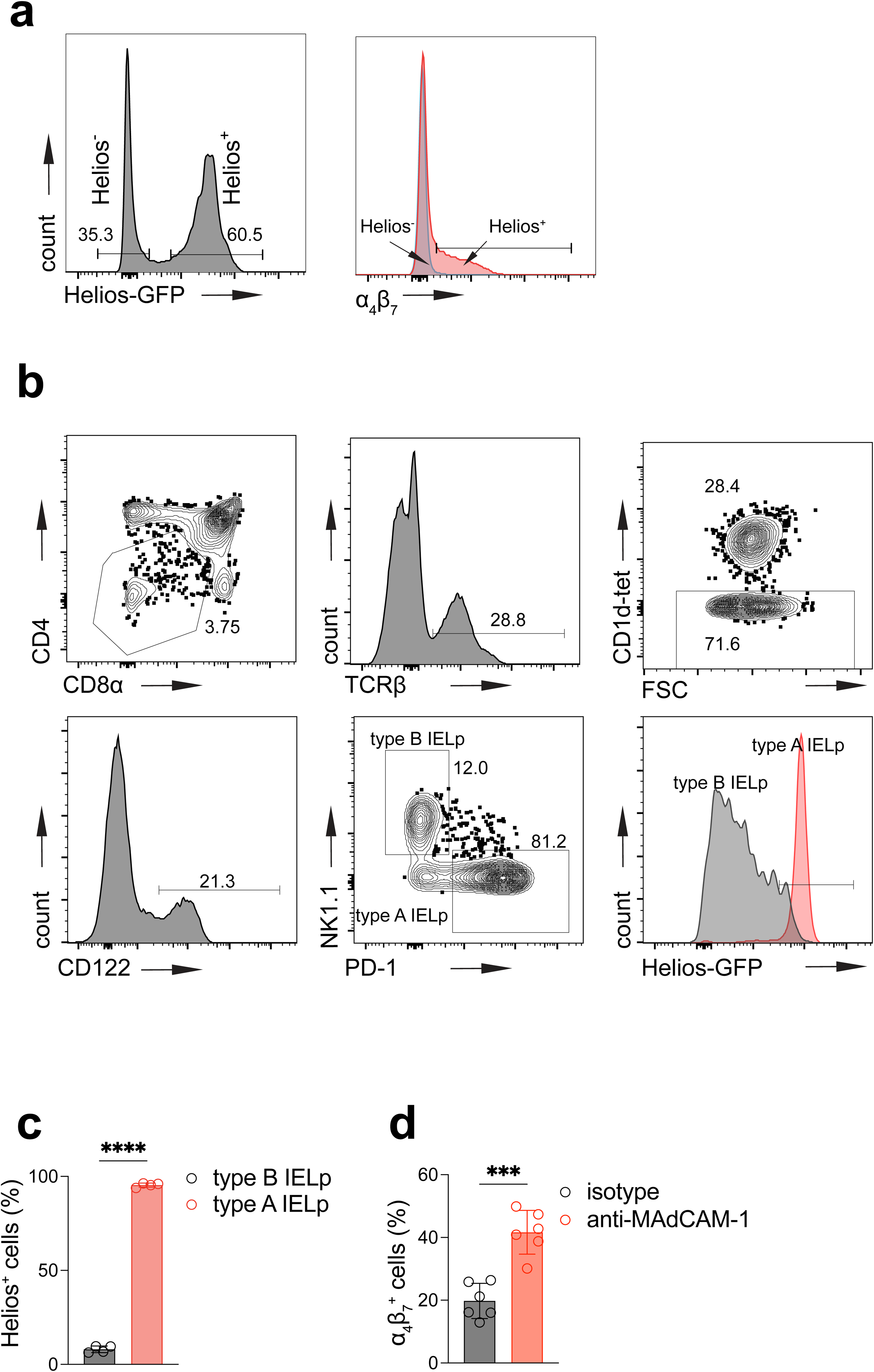
Connections between splenic Helios+CD122+ T cells and natural IELs. **a,** Left, total splenic Ly49+ T cells were gated as either Helios+ or Helios-. Right, representative flow plot showing the expression of α4β7 by each subset, quantified in Fig. 2b. **b,** Gating strategy to identify thymic IELp. CD4-CD8α- (DN) thymocytes are further gated as TCRβ+ (in some experiments, CD5 was used in combination with TCRβ). iNKT cells are excluded from analysis using CD1d tetramer loaded with PBS-57. Mature IELp are identified on the basis of CD122 expression, and are then divided into type A (PD-1+) and type B (NK1.1+) IELp. A representative flow plot shows Helios expression for type A and type B IELp, quantified in **c** (n=4 mice). **d,** As described in Fig. 2f, neonatal mice received anti-MAdCAM-1 blocking antibodies, and were then left untreated for an additional 8-9 weeks. Frequency of α4β7+ cells among splenic Helios+CD122+ T cells is shown (n=6 mice per group). Statistical significance was calculated using Welch’s T test (**c,d**); *** P<0.001, **** P<0.0001. Data are presented as mean values ± SD.

**Extended Data Figure 4:**
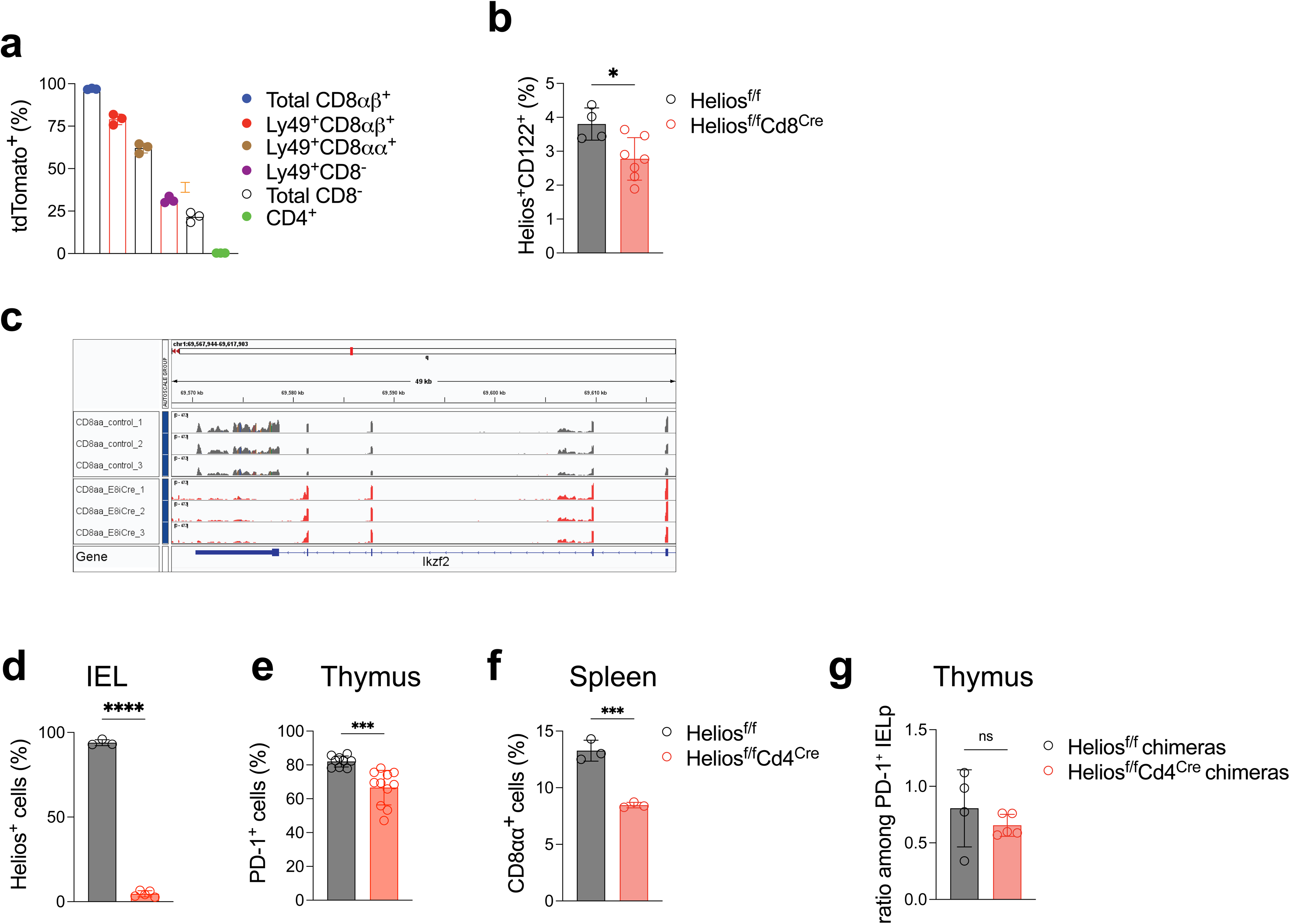
Helios perturbations with Cd8^Cre^, Cd4^Cre^, and in mixed bone marrow chimeras. **a,** Cd8^Cre^ mice were crossed with Ai14 mice (lox-STOP-lox-tdTomato reporter mice) to examine Cre activity among splenic Ly49+ T cells. CD8αβ+ T cells, CD4+ T cells, and CD4-CD8- (“Total CD8-“) T cells were analyzed as control populations, and Ly49+ T cells were divided for analysis based on CD8 expression (n=3 mice). **b,** Frequency of Helios+CD122+ cells among splenic TCRβ+CD4- T cells (n=4, Helios^f/f^; n=7, Helios^f/f^Cd8^Cre^). **c,** IGV tracks generated using BAM files from the bulk RNAseq data as described in Fig. 3b. The leftmost exon displayed is exon 8, which is the floxed exon in Helios^f/f^ mice. **d,** Frequency of Helios+ cells among CD8β- IELs (n=3, Helios^f/f^; n=5, Helios^f/f^Cd4^Cre^). **e,** Frequency of type A IELp (PD-1+ cells) among mature thymic IELp (n=9, Helios^f/f^; n=11, Helios^f/f^Cd4^Cre^). **f,** Frequency of CD8αα+ cells among splenic Ly49+ T cells from Helios^f/f^ and Helios^f/f^Cd4^Cre^ mice (n=3 mice per group). **g,** Ratio of CD45.2:CD45.1/2 cells showing chimerism among thymic type A IELp from mixed bone marrow chimeras. Statistical significance was calculated using Welch’s T test (**d,e,g**) and unpaired T test (**b,f**); ns P≥0.05, * P<0.05, *** P<0.001, **** P<0.0001. Data are presented as mean values ± SD.

**Extended Data Figure 5:**
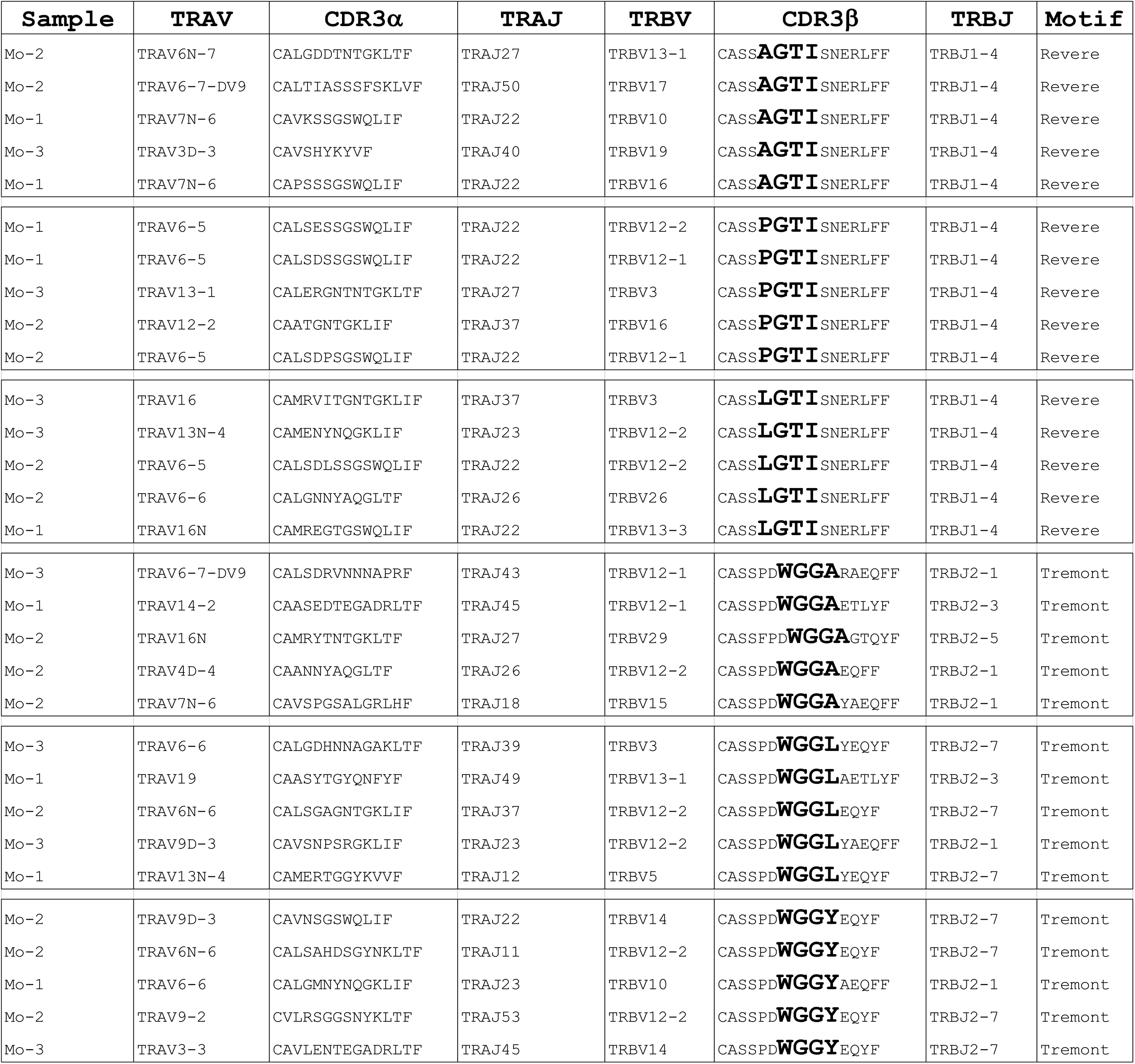
Tables of motif-bearing TCRs. A table shows a selection of TCRs bearing variants of the Revere and Tremont CDR3β motifs.

**Extended Data Figure 6:**
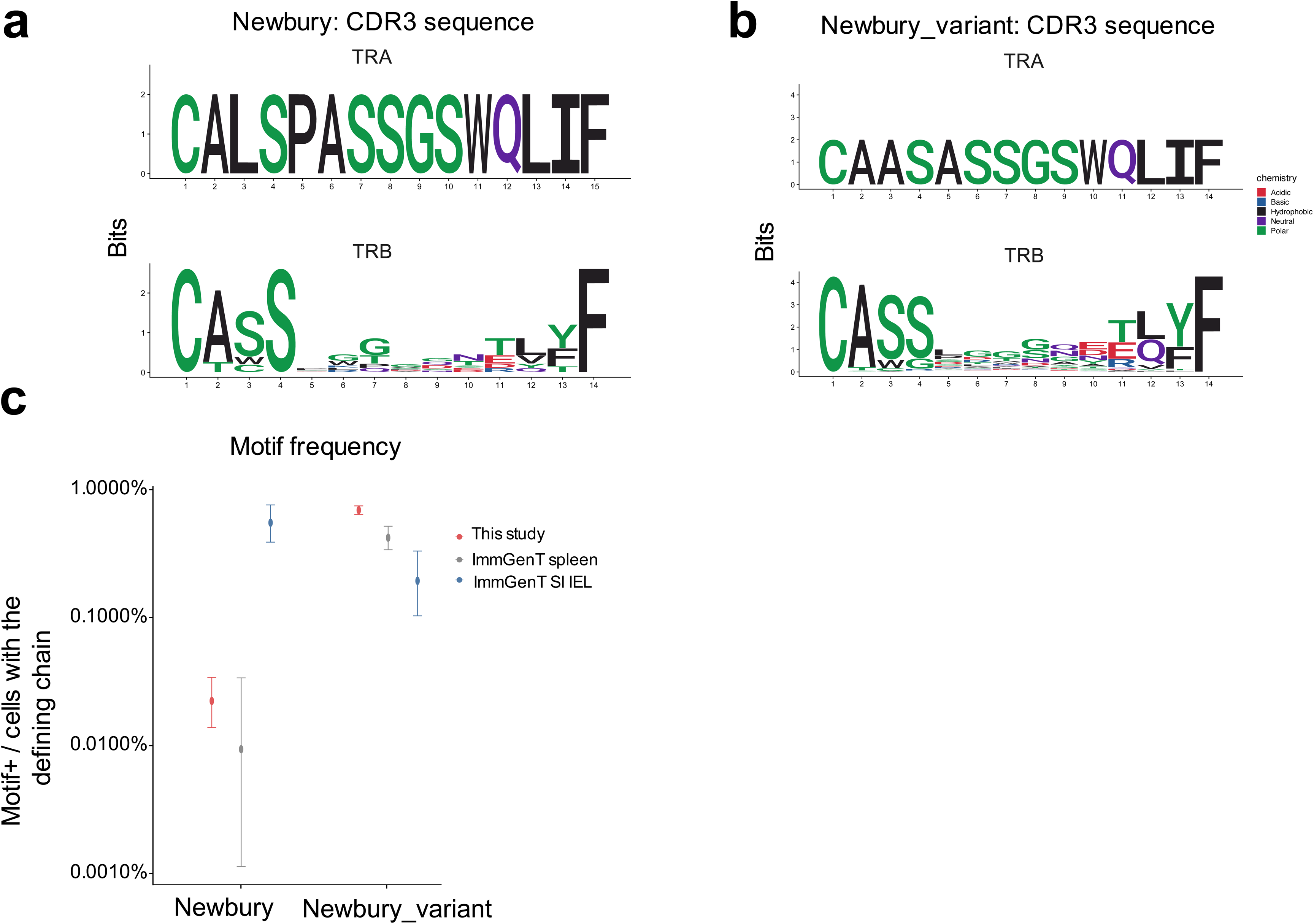
Additional motifs identified in scTCRseq data. **a,** Logos plots showing the definition of a previously identified mammary gland IEL-associated CDR3α sequence motif. We defined the Newbury motif by CDR3α amino acids ‘CALSPASSGSWQLIF’, with no constraint placed on the CDR3β. **b,** Logos plots showing the definition of a similar-appearing CDR3α sequence motif, defined by amino acids ‘CAASASSGSWQLIF’, which we termed ‘Newbury-variant’. **c,** Frequency of Newbury and Newbury-variant motifs in our splenic Ly49+ T cell TCR data (“This study”), ImmGenT data for splenic CD8 T cells, and ImmGenT data for IELs from the small intestine.

**Extended Data Figure 7:**
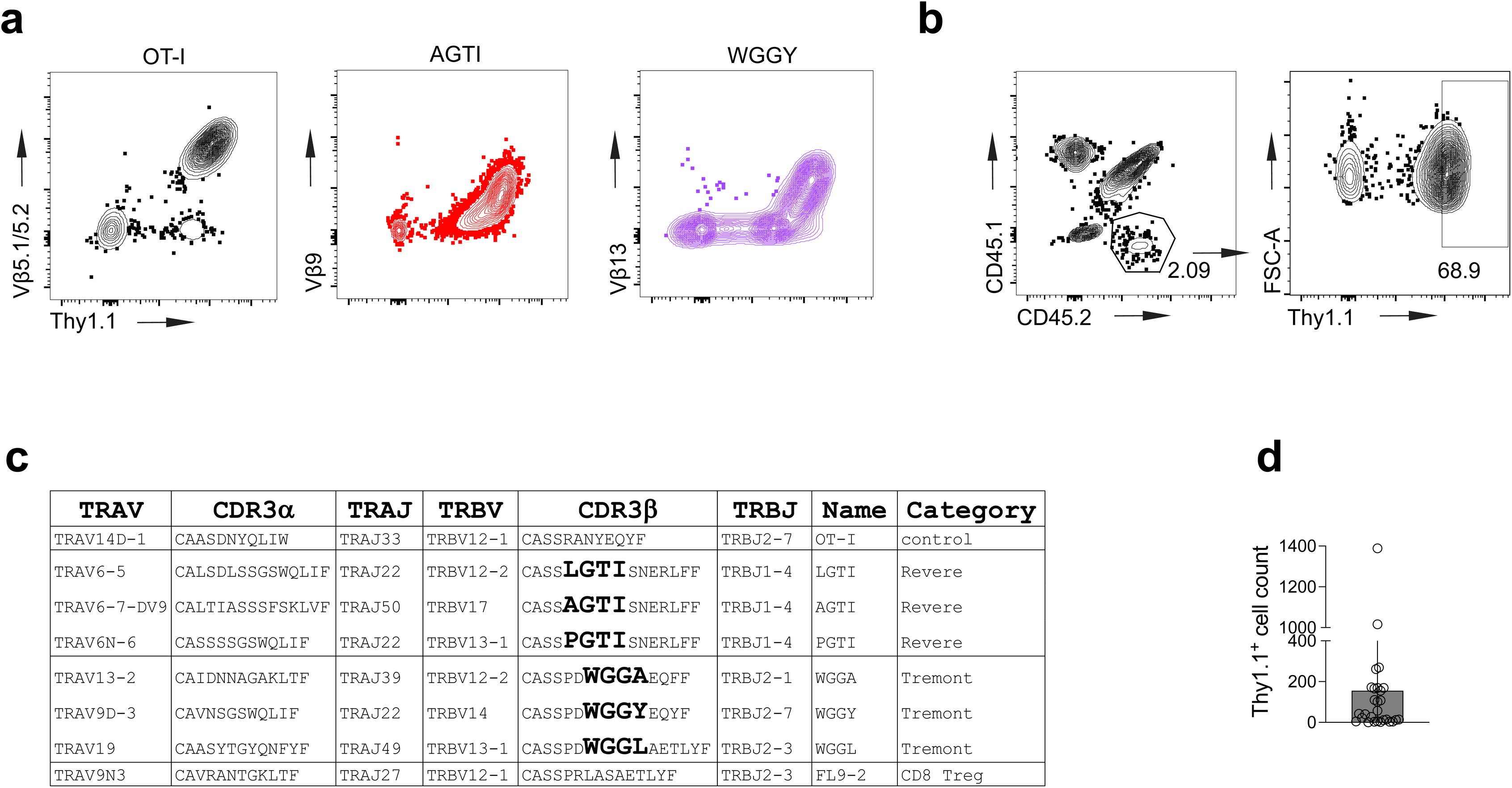
Additional information for TCR retrogenic mice. **a,** Representative flow plots showing staining with different Vβ antibodies to detect expression of retrogenic TCRs. **b,** Example gating strategy to analyze TCRrg cells. Cells are gated as congenic (usually CD45.2+), Thy1.1+ cells. **c,** Table of TCRs tested in TCRrg mice. **d,** Number of Thy1.1+ cells detected in the thymi of TCRrg mice. Data are pooled from 7 batches of TCRrg mice, n=28 mice in total (represents one attempt each for 7 different TCRs). We could not detect adequate numbers of retrogenic thymocytes to evaluate their phenotype.

## Materials and Methods

### Mice

C57BL/6J (B6, strain no. 000664), C57BL/6J-Ptprcem6Lutzy/J (coisogenic CD45.1, strain no. 033076), C57BL/6J-Rag1em10Lutzy/J (coisogenic RagKO, strain no. 034159), C57BL/6-Tg(Cd8a-cre)1Itan/J (Cd8a-Cre, strain no. 008766), B6.Cg-Tg(Cd4-cre)1Cwi/BfluJ (Cd4-Cre, strain no. 022071), and B6.Cg-Gt(ROSA)26Sortm14(CAG-tdTomato)Hze/J (Ai14, strain no. 007914) mice were obtained from the Jackson Laboratory. Helios-GFP and Helios-floxed mice were a kind gift from Ethan Shevach and Angela Thornton. Experimental animals were generated by intercrossing the respective strains. All experimental procedures were performed following the guidelines listed in animal protocol 25-0207, approved by the Institutional Animal Care and Use Committee at Washington University in St. Louis.

### Preparation of cell suspensions

Spleens and thymi were dissected and then mashed through 70um filters in RPMI supplemented with 10% FBS and 1% penicillin and streptomycin (Gibco). ACK lysis was performed to lyse red blood cells, and single cell suspensions were used for downstream applications. For IELs, small intestines were dissected and kept on ice in HBSS (without Ca2^+^ and Mg2^+^) supplemented with HEPES (15mM final concentration). Fecal contents were flushed and Peyer’s patches and residual fat were removed. Intestines were cut open longitudinally, washed vigorously in HBSS/HEPES to remove residual mucous, and cut into ∼1cm-long pieces. Pieces of intestine were incubated in 20mL HEPES/HBSS with 10% FBS and 0.15mg/mL DTT on a revolver/rotator at 24rpm in a 37°C incubator. After 20min, samples were vortexed for 20sec and poured through 100um filters. Cell suspensions were pelleted, resuspended in 40% Percoll, overlaid on 80% Percoll, and spun at 2000rpm for 20min. Cells were taken from the interface and were washed once with PBS before use in downstream applications.

### Flow cytometry

Single cell suspensions were stained with TruStain FcX PLUS (anti-mouse CD16/32) Antibody (BioLegend) and fluorophore-conjugated monoclonal antibodies at 4°C. For intracellular staining to detect transcription factors, the eBioscience Foxp3 / Transcription Factor Staining Buffer Set (Invitrogen) was used; cells were fixed at 4°C and stained at room temperature. Flow cytometry data was collected using a Cytek Aurora spectral flow cytometer, and cell sorting was performed using a BD FACSAriaIII cell sorter.

### Splenic T cell transfers

For splenic T cell transfers, spleens from donor mice were processed (see above), and cells of the same genotype were pooled. Typically, 4 or more donor mice were used for each recipient of Helios+CD122+ T cells. Negative selection using biotinylated antibodies and Streptavidin Microbubbles (Akadeum Life Sciences) was performed prior to sorting, following the manufacturer’s protocol with some modifications. Biotinylated antibodies (all from BioLegend) against CD19 (MB19-1) and CD4 (RM4-5) were added at 31.25ug/100 million cells, and biotinylated antibodies against F4/80 (BM8), CD14 (Sa14-2), Ly6G (1A8), IgM (RMM-1), IgD (11-26c.2a), IgG (Poly4053), TCRgd (GL3), and NKp46 (29A1.4) were added at 6.25ug/100 million cells. Cells were stained at 4°C for 20 minutes. After washing, cells were resuspended and distributed to 5mL Eppendorf tubes, with 50-80 million cells in 500uL separation buffer per tube. 1mL of microbubbles was added per tube, cells and microbubbles were triturated ∼20 times to mix, 3mL separation buffer was added, and then samples were spun at 400g for 5min to separate. After aspirating microbubbles, cells were again pooled according to genotype, and biotinylated CCR9 antibody (CW-1.2) was added at 31.25ug/100 million starting cells. We found that CCR9 was perfectly anti-correlated with CD122 for CD8 T cells (not shown), so we used CCR9 to further enrich for CD122+ T cells by depleting naïve CD8 T cells. After washing, cells were again split to 1-3 5mL Eppendorfs and were depleted with 1mL microbubbles.

After negative selection, cells were stained for sorting with LIVE/DEAD Fixable viability dye (Aqua or NIR, Invitrogen), and then antibodies against CD4 (RM4-4), TCRβ (H57-597), and CD122 (TM-β1). Sorted cells were gated as lymphocytes, singlets (based on FSC and SSC), live, TCRβ+CD4-, and then either Helios+CD122+ or Helios-CD122+. For congenic splenocyte transfers, Helios-GFP mice were used for donors of Helios+CD122+ splenic T cells (CD45.2). The F1 generation of Helios-GFP mice crossed to CD45.1 mice were used for donors of Helios-CD122+ splenic T cells (CD45.1/2). After sorting, cells were mixed and injected retro-orbitally into CD45.1/RagKO recipients. 3-5 weeks later, recipient mice were euthanized, and spleens and IELs were collected for flow cytometry.

To evaluate the ability of splenic T cells to protect mice from DSS-induced colitis, Helios+CD122+ and Helios-CD122+ T cells were simultaneously sorted from Helios-GFP mice. Sorted cells were expanded in 100ng/mL IL-15 (BioLegend) for 7 days before cell transfer. Cultured cells (3-6 million per recipient in PBS) were transferred to recipient CD45.1/RagKO mice via retro-orbital injection 4 days before induction of DSS colitis (PBS alone was injected into control animals).

### In vitro culture of sorted Ly49+ T cells with different concentrations of IL-15

Spleens and lymph nodes from Helios-GFP mice were processed, and negative selection was performed using Akadeum microbubbles, following largely the same protocol as for splenic T cell transfers (see above). A second round of depletion with CCR9 was not performed. After negative selection, cells were stained with LIVE/DEAD Fixable viability dye (Aqua), and then antibodies against CD3 (17A2), CD8α (53-6.7), CD122 (TM-β1), CD44 (IM7), CD8β (YTS156.7.7), and Ly49 family members (Ly49F, HBF-719; Ly49G, 4D11; Ly49A, A1; Ly49I, YLI-90; Ly49H, 3D10; Ly49D, 4E5; Ly49E/F, CM4; Ly49C/I, 5E6; Ly49FCIH, 14B11; all on the same fluorophore). Cells were gated as lymphocytes, singlets (based on FSC and SSC), Live, CD3+CD8α+, CD44+CD122+, Ly49+, and then either Helios-, Helios+CD8αβ+, or Helios+CD8αα+. Each sorted population was split into two wells and cultured in either 10ng/mL IL-15 or 100ng/mL IL-15 (BioLegend). Cells were initially cultured in a 96-well U-bottom plate and were expanded to a 12-well plate and split as needed.

### Antibody treatment

Helios-GFP mice were injected i.p. with 200ug of anti-α4β7 (clone DATK32, Leinco), anti-MAdCAM-1 (clone MECA-367, Leinco), or isotype control (clone 1-1, Leinco) antibody every 3 days, for a total of 5 doses. For dual blockade, mice were given 200ug of both blocking antibodies, and the isotype control mice received 400ug of isotype control antibody. For neonatal integrin blockade, male and female pups were distributed evenly between treatment groups, and the first dose was given on day 11 of life.

### DSS colitis

DSS colitis was induced according to previously established protocols with minor modifications^50^. Mice were administered 3% DSS (TdB Labs) in their drinking water for 8-10 days. Body weight was measured daily, and mice were euthanized if they reached less than 80% of initial weight.

### Generation of mixed bone marrow chimeric mice

Recipient mice (CD45.1, 12 weeks old) were lethally irradiated with 1000 cGy. Femurs and tibiae from donor mice were dissected and cleaned of tissue, and bone marrow cells were isolated by centrifugation at 10,000 x g for 15 seconds, as previously described with minor modifications^51^. Cells were passed through 70um filters and ACK lysis was performed to remove red blood cells. Cells were counted, and 10 million cells of each genotype per recipient were pooled. Marrow from Helios^f/f^Cd4^Cre^ donors was mixed with CD45.1/2 marrow to generate Helios^f/f^Cd4^Cre^ chimeras; marrow from Helios^f/f^ littermates was mixed with CD45.1/2 marrow to generate Helios^f/f^ chimeras. Bone marrow cells were resuspended in 100uL of PBS per recipient and were injected retro-orbitally. Recipients were started on antibiotics (80mg/L trimethoprim and 400mg/L sulfamethoxazole, in drinking water) 3 days before irradiation and were maintained on antibiotics for two weeks. Spleens, thymi, and IELs from recipient mice were analyzed >12 weeks after transplantation.

### TCR construct design

The timed TCR expression construct was modified from an existing conditional retroviral vector, pMGflThy1.1, which was a kind gift of P. Savage^52^. That vector contains a GFP sequence in a loxP-flanked STOP cassette, upstream of a TCRα coding segment. We replaced the GFP sequence with an ‘irrelevant’ TCRβ chain, which facilitates the passage of beta selection by developing thymocytes, and is then removed by Cd4^Cre^-mediated excision. We used the TCRβ chain from the OT-I TCR, and obtained the sequence from Addgene plasmid #52111 (OTI-2A.pMIG II). We then replaced the downstream TCRα coding segment with linked TCRα-2A-TCRβ sequences taken from paired TCRs of interest. We identified TCRs of interest in our scTCRseq data (see below). To generate each full length TCR, we combined the CDR3 region from our sequencing data with sequences for that TCR’s V- and J-genes; V- and J-gene sequences (including the leader sequence) were obtained from IMGT. Sequences were further codon-optimized before being synthesized and cloned into the vector. Gene synthesis and cloning was performed by GenScript, and plasmid sequences were confirmed by plasmid sequencing via Plasmidsaurus.

### Retrovirus production

Retrovirus was produced using Plat-E packaging cells (Cell Biolabs). Conditional TCR constructs were transfected into Plat-E cells using Lipofectamine LTX Reagent with PLUS Reagent (Invitrogen), and viral supernatants were harvested after 48 and 72 hours. Viral supernatants were filtered through 0.45um filters, concentrated using Retro-X Concentrator (Takara), snap-frozen in liquid nitrogen, and stored at −80°C until use.

### TCR retrogenic mouse generation

Donor Cd4^Cre^RagKO mice were injected i.p. with 3mg of 5-fluorouracil (Fresenius Kabi) 3 days prior to bone marrow isolation. Bone marrow was isolated by centrifugation (see above, ‘Generation of mixed bone marrow chimeras’). After ACK lysis, hematopoietic progenitor cells (HPCs) were enriched using magnetic negative selection (EasySep Mouse Hematopoietic Progenitor Cell Isolation Kit, Stemcell Technologies). HPCs were cultured in X-Vivo 10 medium (Lonza) supplemented with 15% FBS and 1% penicillin and streptomycin (Gibco). The media was further supplemented with 10ng/mL mouse IL-6, 6ng/mL mouse IL-3, 100ng/mL mouse SCF, and 50ng/mL mouse Flt3L (all from PeproTech). Cultures were maintained at 37°C with 5% CO2. Media was exchanged after 1 day in culture, and cells were typically split 1:2 as well.

After 2 days in culture, HPCs were transduced with TCR-encoding retroviruses using RetroNectin (Takara). 6-well plates were coated with RetroNectin at 4°C overnight; 20ug RetroNectin in 1.5mL PBS was used per well, and typically two wells were used per TCR. The “RetroNectin-Bound Virus” method was used for transduction. Thawed retroviral supernatant was added to RetroNectin-coated plates and spun at 2500g for 90 min in a 32°C centrifuge. HPCs (typically 500-800k cells per well) were then added to the plates and spun at 500g for 10 min to settle the cells onto the virus. The next day, a second round of transduction was performed. Cells were typically split the day after the second transduction and were transplanted after a total of 4-5 days in culture. On the day of transplantation, recipient mice (CD45.1) were irradiated with 1000 cGy. Freshly harvested “filler” bone marrow cells were taken from congenically distinct (typically CD45.1/2) WT donors on the day of transplantation. We found that 1 million filler cells were sufficient to generate low frequency retrogenic mice, and 1 million filler cells were used per recipient in all retrogenic mouse experiments. Mixed filler marrow and transduced HPCs were injected retro-orbitally into the irradiated recipients. The recipient mice were started on antibiotics 5 days before irradiation and were maintained on antibiotics for 2 weeks after irradiation. Roughly 4 weeks after transplantation, cheek bleeds were performed to confirm expression of transduced TCRs by flow cytometry of peripheral blood. Flow cytometry was performed on spleens and IELs >8 weeks after reconstitution.

### Bulk RNAseq of CD8αα+ IELs

IELs were isolated from the small intestines of Helios^f/f^Cd8^Cre^ mice and Helios^f/f^ littermates (n=3 per genotype, 8-10 weeks old). After Percoll density gradient, cells were stained with LIVE/DEAD Fixable viability dye (Aqua), and then antibodies against CD45 (30-F11), CD3 (17A2), CD4 (RM4-5), TCRβ (H57-597), CD8α (53-6.7), and CD8β (YTS156.7.7). Cells were gated as lymphocytes, singlets (based on FSC and SSC), live and CD45+, CD3+, TCRβ+CD4-, and then CD8α+CD8β-. CD8αα+ IELs were sorted directly into Buffer RLT Plus (Qiagen) with 2-mercaptoethanol (Calbiochem), vortexed for 1min., and placed on dry ice. Once all samples were sorted, the lysates were stored at −80°C. RNA was extracted using the RNeasy Plus Micro kit (Qiagen).

Library preparation and sequencing was performed by the Genome Technology Access Center at the McDonnell Genome Institute. Total RNA integrity was determined using Agilent Bioanalyzer or 4200 Tapestation. Library preparation was performed with 10ng of total RNA with a Bioanalyzer RIN score greater than 8.0. ds-cDNA was prepared using the SMARTer Ultra Low RNA kit for Illumina Sequencing (Takara-Clontech) per manufacturer’s protocol. cDNA was fragmented using a Covaris E220 sonicator using peak incident power 18, duty factor 20%, cycles per burst 50 for 120 seconds. cDNA was blunt ended, had an A base added to the 3’ ends, and then had Illumina sequencing adapters ligated to the ends. Ligated fragments were then amplified for 12-15 cycles using primers incorporating unique dual index tags. Fragments were sequenced on an Illumina NovaSeq X Plus using paired end reads extending 150 bases. Basecalls and demultiplexing were performed with Illumina’s bcl2fastq software with a maximum of one mismatch in the indexing read. RNA-seq reads were then aligned to the Ensembl GRCm39.113 primary assembly with STAR version 2.7.11b. Gene counts were derived from the number of uniquely aligned unambiguous fragments by Subread:featureCount version 2.0.8. Differential gene expression analysis was performed using the limma package^53^.

### scRNAseq and scTCRseq of splenic Ly49+ T cells

Splenocytes were isolated from B6 mice (n=3) and negative selection was performed using biotinylated antibodies (all from BioLegend) against CD19 (MB19-1), CD4 (RM4-5), F4/80 (BM8), IgM (RMM-1), IgD (11-26c.2a), and IgG (Poly4053), and Streptavidin Microbubbles (Akadeum Life Sciences). Cells were then stained with TruStain FcX PLUS (anti-mouse CD16/32) Antibody (BioLegend), LIVE/DEAD Fixable viability dye (Aqua), and then antibodies against CD19 (6D5), CD3 (17A2), and Ly49 family members (Ly49F, HBF-719; Ly49G, 4D11; Ly49A, A1; Ly49I, YLI-90; Ly49H, 3D10; Ly49D, 4E5; Ly49E/F, CM4; Ly49C, 4LO3311; Ly49C/I, 5E6; Ly49FCIH, 14B11; all on the same fluorophore). Cells were also stained with TotalSeq antibodies (all from BioLegend) against TCRβ (C0120), TCRγδ (C0986), CD4 (C0001), CD8α (C0002), CD8β (C0230), CD44 (C0073), CD122 (C0227), NK1.1 (C0118) and TotalSeq-C hashtag antibodies (a different hashtag was used for each sample) for CITE-seq analysis. Sorted cells were gated as lymphocytes, singlets (based on FSC and SSC), live, CD3+CD19-, and Ly49+ cells. Samples were pooled for encapsulation (10X Chromium X). Libraries were prepared using Chromium Next GEM Single Cell 5’ HT Kit v2 reagents and were sequenced on a Novaseq X Plus.

### scRNAseq alignment and processing

Reads were processed per library with Cell Ranger multi (Cell Ranger v7.2.0, 10x Genomics). Gene expression reads were aligned to the mm10 mouse reference (refdata-gex-mm10-2020-A). Antibody capture reads were counted against a custom feature reference. TCR contigs were assembled and annotated using the refdata-cellranger-vdj-GRCm38-atls-ensembl-5.0.0. Per-library filtered feature and contig annotation outputs were carried forward. Hashtag counts were normalized using centered log-ratio and demultiplexed with HTODemux in Seurat v5.3.1.0^54^. Cells were retained with 800 to 6,000 detected genes, under 10% mitochondrial reads, 200 to 10,000 antibody-derived tag counts, and 300 to 12,000 hashtag counts. Mitochondrial content used the “mt-“ pattern and ribosomal content used “Rp[sl][0-9]|Rplp[0-9]|Rpsa” pattern.

### scRNAseq normalization, integration and clustering

RNA counts were log-normalized using a scale factor of 10,000. The top 3,000 variable features were selected by variance-stabilizing transformation. Adaptive immune receptor gene segments, ribosomal, mitochondrial, and hemoglobin genes were removed from the variable gene set. The retained features were scaled, and a principal component analysis was run with downstream analysis using the first 30 dimensions. Lane-level batch effects were corrected with Harmony (v1.2.4) on the PCA embeddings using θ = 2, λ = 1, and a maximum of 20 iterations^55^. A shared nearest neighbor graph was built from the Harmony-corrected 30 dimensions. Leiden clusters were calculated across a resolution of 0.2 to 1.0, with a resolution of 0.6 selected for optimal silhouette scoring. UMAP coordinates were computed on the same 30 Harmony dimensions using uwot v0.2.4. One cluster was manually excluded due to an abnormally low count and a feature space consistent with technical noise. Cluster markers were identified using a Wilcoxon rank-sum test on an object downsampled to 400 cells per cluster, restricting to positive markers with a minimum detection fraction of 0.25 and a minimum log2 fold-change of 0.25. Module scores for curated gene sets were computed with Seurat. The Pearson correlation between *Cd8b1* and *Ikzf2* was computed using pseudo-bulking of the raw counts by cluster-animal groupings, with groups of fewer than 20 cells excluded. PLZF+ innate-like and terminal effector clusters were manually excluded. A random-intercept model using lmerTest v3.1.3 on lme4 v1.1.38 was used to calculate the slope^56^.

### TCR repertoire assembly and analysis

Per-lane filtered contigs were processed using scRepertoire v2.5.7^57^, retaining the highest-read contigs for the alpha and beta TCR chains. Motif families were defined using the following scheme, following previous work that identified the motifs^33^: 1) Revere, TRB chain, “CASS.GTISNERLFF” using the TRBJ1-4 gene segment; 2) Tremont, TRB chain “CASSPDWGG[A-Z]{3,8}F”; 3) Newbury, TRA chain, “CALSPASSGSWQLIF”; and 4) Newbury variant, TRA chain, “CAASASSGSWQLIF”. The external reference table was built from the ImmGenT per-donor tables, using only CD8+ T cells from wild-type donors^58^. ImmGenT data IMGT nomenclature was harmonized with the Cell Ranger outputs, removing allele designation, D and N copies, and TRAv6-7/DV9-style naming. Generation probability was calculated using the OLGA v1.3.0 model^59^ from immLynx v1.0.0^60^ using the bundled VJ alpha and VDJ beta mouse models, with TCR sequences scoring 0 removed (n=195). Motif frequencies were calculated using the per-chain denominators with exact 95% Clopper-Pearson binomial intervals and using Fisher exact test with Benjamini-Hochberg correction. Per-family alluvial diagrams were drawn with ggsankey v0.0.99999. CDR3 sequence logos were drawn with ggseglogo v0.2 using bits.

## Acknowledgements

We would like to thank Brian Edelson, Takeshi Egawa, and Gwendalyn Randolph for critical feedback on the project and manuscript. We would also like to thank the Genome Technology Access Center at the McDonnell Genome Institute for their sequencing services and Department of Pathology and Immunology flow cytometry core for their assistance with flow cytometry and cell sorting. We would like to thank Stephen Jameson and Ka Hyun Rhee for critical discussion and input on the manuscript. Some figures were created with BioRender.com.

## Author contributions

Conceptualization: S.J.H.M., N.S.

Data curation: S.J.H.M., N.B., N.S.

Investigation: S.J.H.M., N.S.

Methodology: S.J.H.M., T.Y., C.T., J.B., N.B., N.S.

Resources: S.J.H.M., N.B., W.M.Y., N.S.

Writing (Initial draft): S.J.H.M., N.S.

Writing (Review and editing): all authors reviewed the manuscript

Supervision: N.S.

## Data and materials availability

Sequencing datasets (bulk RNAseq and scRNA+TCRseq) are being submitted to Genome Expression Omnibus (GEO) and accession numbers will be added once they are received. Requests for further information or access to data should be directed to and will be fulfilled by the corresponding author, Naresha Saligrama.

